# Metabolic flux partitioning between the TCA cycle and glyoxylate shunt combined with a reversible methyl citrate cycle provide nutritional flexibility for *Mycobacterium tuberculosis*

**DOI:** 10.1101/2021.01.29.428863

**Authors:** Khushboo Borah, Tom A. Mendum, Nathaniel D. Hawkins, Jane L. Ward, Michael H. Beale, Gerald Larrouy-Maumus, Apoorva Bhatt, Martine Moulin, Michael Haertlein, Gernot Strohmeier, Harald Pichler, V. Trevor Forsyth, Stephen Noack, Celia W. Goulding, Johnjoe McFadden, Dany J.V. Beste

**Affiliations:** Department of Microbial and Cellular Sciences, Faculty of Health and Medical Sciences, University of Surrey, Guildford GU2 7XH, UK; Department of Computational and Analytical Sciences, Rothamsted Research, Harpenden, Herts AL5 2JQ, UK; MRC Centre for Molecular Bacteriology and Infection, Department of Life Sciences, Faculty of Natural Sciences, Imperial College London, London, SW7 2AZ, UK; School of Biosciences, University of Birmingham, Edgbaston, Birmingham, B15 2TT, UK; Life Sciences Group, Institut Laue-Langevin, 71 avenue des Martyrs CS 20156, 38042 Grenoble Cedex 9, France; Partnership for Structural Biology, 71 avenue des Martyrs CS 20156, 38042 Grenoble Cedex 9, France; Austrian Centre of Industrial Biotechnology, Petersgasse 14, 8010 Graz, Austria; Institute of Organic Chemistry, NAWI Graz, Graz University of Technology, Stremayrgasse 9, 8010 Graz, Austria; Institute of Molecular Biotechnology, NAWI Graz, BioTechMed Graz, Graz University of Technology, Petersgasse 14, 8010 Graz, Austria; Faculty of Natural Sciences, Keele University, Staffordshire ST5 5BG, UK; Forschungszentrum Jülich GmbH, Institute of Bio- and Geosciences 1: Biotechnology 2, Jülich, Germany; Department of Pharmaceutical Sciences & Molecular Biology & Biochemistry, University of California Irvine, Irvine, CA 92697, US

**Keywords:** Tuberculosis, metabolism, metabolic flux, chemostat, mass isotopomer, tri-carboxylic acid cycle, steady state, cholesterol, intracellular

## Abstract

The utilisation of multiple host-derived carbon substrates is required by *Mycobacterium tuberculosis* (Mtb) to successfully sustain a tuberculosis infection thereby identifying the Mtb specific metabolic pathways and enzymes required for carbon co-metabolism as potential drug targets. Metabolic flux represents the final integrative outcome of many different levels of cellular regulation that contribute to the flow of metabolites through the metabolic network. It is therefore critical that we have an in-depth understanding of the rewiring of metabolic fluxes in different conditions. Here, we employed ^13^C-metabolic flux analysis using stable isotope tracers (^13^C and ^2^H) and lipid fingerprinting to investigate the metabolic network of Mtb growing slowly on physiologically relevant carbon sources in a steady state chemostat. We demonstrate that Mtb is able to efficiently co-metabolise combinations of either cholesterol or glycerol along with C2 generating carbon substrates. The uniform assimilation of the carbon sources by Mtb throughout the network indicated no compartmentalization of metabolism in these conditions however there were substrate specific differences in metabolic fluxes. This work identified that partitioning of flux between the TCA cycle and the glyoxylate shunt combined with a reversible methyl citrate cycle as the critical metabolic nodes which underlie the nutritional flexibility of Mtb. These findings provide new insights into the metabolic architecture that affords adaptability of Mtb to divergent carbon substrates.

**Importance:** Each year more than 1 million people die of tuberculosis (TB). Many more are infected but successfully diagnosed and treated with antibiotics, however antibiotic-resistant TB isolates are becoming ever more prevalent and so novel therapies are urgently needed that can effectively kill the causative agent. Mtb specific metabolic pathways have been identified as an important drug target in TB. However the apparent metabolic plasticity of this pathogen presents a major obstacle to efficient targeting of Mtb specific vulnerabilities and therefore it is critical to define the metabolic fluxes that Mtb utilises in different conditions. Here, we used ^13^C-metabolic flux analysis to measure the metabolic fluxes that Mtb uses whilst growing on potential *in vivo* nutrients. Our analysis identified selective use of the metabolic network that included the TCA cycle, glyoxylate shunt and methyl citrate cycle. The metabolic flux phenotypes determined in this study improves our understanding about the co-metabolism of multiple carbon substrates by Mtb identifying a reversible methyl citrate cycle and the glyoxylate shunt as the critical metabolic nodes which underlie the nutritional flexibility of Mtb.

## Introduction

*Mycobacterium tuberculosis* (Mtb) is the causative agent of a global tuberculosis (TB) pandemic which has now reached staggering levels making Mtb once again a leading cause of death globally (1). The terrifying trend of increasing antibiotic resistant TB is destabilising TB control measures making new therapeutics which target drug-resistant strains of Mtb an urgent priority (1–3). Mtb is an unusual bacterial pathogen, which has the remarkable ability to cause both acute life threatening disease and also clinically latent infections that can persist for the lifetime of the human host. Experimental evidence has identified central carbon metabolism as instrumental to this pathogenic strategy and therefore our research is focused on investigating the metabolic capabilities of Mtb both *in vitro* and *ex vivo* (4–7).

Mtb maintains a functional tricarboxylic acid (TCA) cycle, pentose phosphate pathway (PPP) and Embden-Meyerhof-Parnas pathway (EMP), as well as enzymes providing a metabolic link between glycolysis and the TCA cycle (4,8). Mtb also has two alternative pathways (methyl citrate cycle and the B12-dependent methyl malonyl pathway) for metabolizing propionyl-CoA derived from the metabolism of sterols, uneven branched chain fatty acids and amino acids (9–10). These pathways allow Mtb to utilise a wide range of carbon sources that includes carbohydrates, sugars, fatty acids, amino acids and sterols (5, 8, 11–13). Whilst the basic architecture of central carbon metabolism of Mtb is now well established there are still many questions regarding how the flux of metabolites through this network are modulated under various different nutritional conditions.

Using stable isotope labelled nutrients for studying the metabolism of Mtb has proved extremely informative (4, 6, 9, 14–15). Metabolic labelling experiments using ^13^C labelled combinations of acetate, glycerol and glucose have demonstrated that Mtb is able to co-catabolise two carbon sources simultaneously demonstrating that Mtb does not use carbon catabolite repression to regulate metabolism (14). This work also suggested that Mtb not only co-catabolized these carbon substrates but did so in a compartmentalized and segregated manner. However, importantly this work was not able to determine the metabolic flux profiles on these combinations of carbon substrates. Understanding metabolic fluxes of Mtb during co-catabolism of multiple carbon sources will allow us to identify nodes of metabolism most amenable to therapeutic intervention. By combining isotopomer labelling with our steady state chemostat model system of mycobacterial growth allowed us to perform ^13^C-metabolic flux analysis (MFA) at different growth rates in carbon limited conditions (4). Previously we identified the activity of a novel GAS pathway for pyruvate dissimulation when Mtb was growing slowly on glycerol and oleic acid and demonstrated that the pathway requires isocitrate lyase and the enzymes of the anaplerotic node (4) both of which are important for the survival of Mtb in the host (5).

Glycerol was not considered an important carbon source for Mtb as glycerol kinase (*glpK*), which is an essential gene for the conversion of glycerol to glycerol 3-phosphate (16) is dispensable for the growth of Mtb in a murine TB model (17). However, the detection of glycerol in human-like TB lesions has compelled a re-evaluation of the role of glycerol and glycerol containing metabolites in the life cycle of Mtb (18). Moreover, Mtb has been shown to co-metabolise a mixture of carbon substrates when growing in host macrophage cells that included an unknown glycolytic C3 substrate (5) that could potentially be glycerol.

Metabolomic and gene-deletion studies have highlighted fatty acids and cholesterol as critical to the nutrition, survival and virulence of Mtb (11, 16,19–21) and that Mtb encounters and co-metabolises both substrates simultaneously *in vivo* (5, 11). Cholesterol, uneven chain fatty acids and branched chain amino acids are all catabolised to provide Mtb with the metabolite propionyl-CoA which although required for the synthesis of important virulence cell wall lipids, (phthiocerol-dimycocerosate (PDIM), polyacylated trehalose and sulfolipids) is also toxic if allowed to accumulate intracellularly (22–24). In addition to metabolising propionyl CoA through the methyl citrate cycle (MCC) or in the presence of vitamin B12, the methyl malonyl cycle, Mtb can also sequester propionyl CoA into methyl branched cell wall lipids (22, 23). During infection it is thought that Mtb uses fatty acids to prime this process further suggesting that cholesterol and fatty acid metabolism occurs simultaneously *in vivo* (19, 24–26). Despite numerous biochemical studies to elucidate the biochemical degradation pathways and studies exploring the role of specific enzymes in cholesterol and fatty acid metabolism the metabolic flux profile of Mtb growing on this combination of substrates has never been directly measured.

In this study we performed ^13^C-MFA on steady state, slowly growing cultures of Mtb using an extended version of our ^13^C isotopomer model (4), which includes the MCC. We compared the metabolic flux profile of chemostat cultures of Mtb growing in carbon limited conditions growing with cholesterol/acetate with those growing on glycerol/Tween80 (oleic acid). We demonstrate that when Mtb is growing slowly on cholesterol and acetate Mtb utilises a complete TCA cycle in combination with the glyoxylate shunt. There is very little demand for the MCC in these conditions as lipid fingerprinting identified that the propionyl-CoA derived cholesterol is being preferentially incorporated into lipids. Conversely when growing on glycerol and Tween80, Mtb utilises an incomplete TCA cycle and the methyl citrate cycle is reversed to provide propionyl CoA for the synthesis of virulence lipids. This work highlights that re-routing fluxes through the TCA cycle, MCC and the glyoxylate shunt and in particular the ability to alternate the direction of the MCC whilst co-metabolising carbon substrates during slow growth is critical to the metabolic flexibility of Mtb.

## Results

### Metabolic profile of Mtb during steady state growth and co-catabolism of multiple carbon substrates

To define the metabolic profile of Mtb growing on cholesterol and acetate, Mtb H37Rv cultures were grown in carbon limited chemostats operating at a dilution rate of μ=0.01 h (doubling time (t_d_)=69 h). We adopted this dilution rate as we have previously demonstrated that the transcriptional response at this growth rate has many similarities to the transcriptional response characteristic of the adaptation of Mtb to the macrophage environment, and others have shown a similar profile from Mtb isolated from sputum (16). The cultures were grown in Roisin’s minimal medium (4) with a combination of EITHER cholesterol/acetate (CHL-ACE), OR glycerol/Tween 80 (GLY-OLA) as previously described (4). After approximately three volume changes little variation was observed in the CO2 and biomass production rates, indicating that a metabolic steady state conditions had been attained (Fig.S1 in supplementary material). Data from the chemostat cultures demonstrated that Mtb was able to co-metabolise acetate and cholesterol simultaneously with similar yields of bacteria to cultures growing on glycerol and Tween80. Substrate uptake rates were also comparable under the two growth conditions, but the CO2 production rate was significantly higher in CHL-ACE (Table 1).

**Table 1:** Steady state characteristics of *Mycobacterium tuberculosis* grown in carbon limited chemostats.

| Carbon sources | CHL-ACE | GLY-OLA |
| --- | --- | --- |
| Dilution rate ($\text{h}^{-1}$) | 0.01 | 0.01 |
| Biomass (g dry weight $\text{l}^{-1}$) | 0.49 | 2.18 |
| CFU ($\times 10^7 \text{ ml}^{-1}$) | 3.9 | 33 |
| Cell weight (pg dry weight $\text{cfu}^{-1}$) | 12.5 | 6.7 |
| Substrate consumption rate ($\text{mmol g biomass}^{-1} \text{ h}^{-1}$) | Acetate = 0.26<br>Cholesterol = 0.0085 | Glycerol = 0.23<br>Oleic acid = 0.002 |
| $\text{CO}_2$ production rate ($\text{mmol g biomass}^{-1} \text{ h}^{-1}$) | 0.245 | 0.178 |
| Yield ($\text{g biomass}^{-1} \text{ mmol carbon}^{-1}$) | 0.013 | 0.012 |

For the labelling experiments steady state chemostat cultures were switched to identical media containing ^13^C labelled substrates (30% [^13^C_3_]glycerol OR 100% [^13^C_2_]acetate). Samples were taken every volume change for a total of four volume changes to ensure that an isotopic stationary state was attained. Labelling of proteinogenic amino acids and intracellular metabolites were measured using gas chromatography-mass spectrometry (GC-MS) or liquid chromatography-mass spectrometry (LC-MS) as previously described (4). The labelling pattern of the amino acids changed very little between the third and fourth volume change confirming that the culture had reached an isotopic steady state which is essential for determining intracellular fluxes using ^13^C-MFA (Table 1; Fig. S1C, D).

Previous studies indicated that Mtb was operating a form of compartmentalised metabolism when growing on solid 7H10 agar containing combinations of the carbon sources glucose, glycerol and acetate whereby individual substrates had distinct metabolic fates (14). However our ^13^C labelling data from chemostat cultures at metabolic and isotopic steady state showed no evidence of such compartmentalisation when Mtb was grown with either GLY-OLA or CHL-ACE as evidenced by uniform distribution of both unlabelled and ^13^C labelled substrates (Fig. 1).

**Fig.1.**
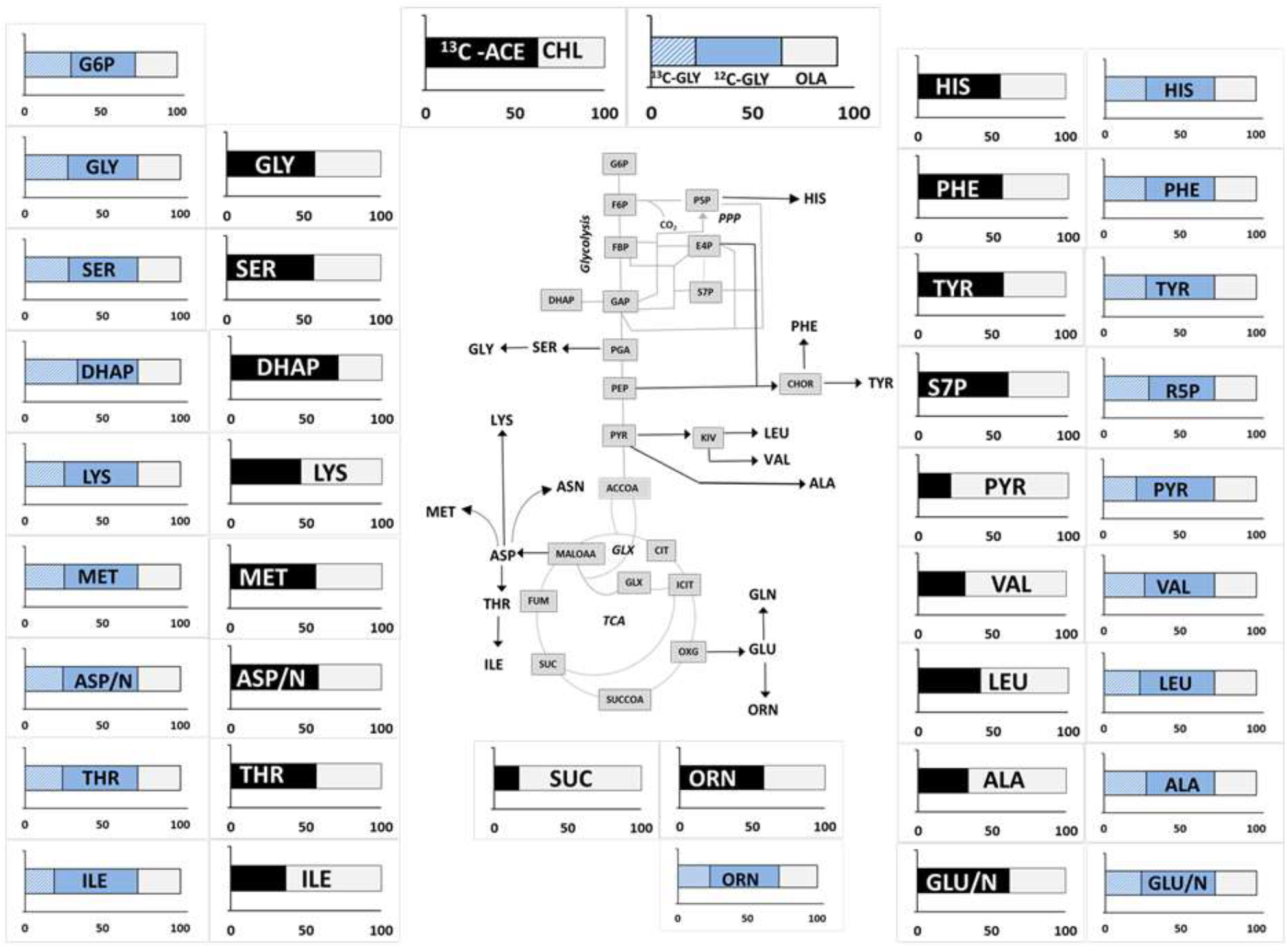
^13^C incorporation into the proteinogenic amino acids and intracellular metabolites from metabolic and isotopic steady state chemostat Mtb cultures. Label distribution is shown in metabolites from Mtb grown in 30% [^13^C_3_] glycerol and unlabelled Tween80 which provides Mtb with oleic acid (GLY-OLA) and 100% [^13^C_2_] acetate and unlabelled cholesterol (CHL-ACE). Average ^13^C incorporation was calculated for metabolites harvested at a steady state growth and are shown as the amount labelled (^13^C) and unlabelled (^12^C). Data are average of 3 replicate measurements and shown for DHAP (dihydroxyacetone phosphate), PYR (pyruvate) ALA (alanine), GLY (glycine), SER (serine), LYS (lysine), MET (methionine), ASP/N (aspartate/asparagine), THR (threonine), ILE (isoleucine), ORN (ornithine), GLU/N (glutamate/glutamine), SUC (succinate), LEU (leucine), ILE (isoleucine), VAL (valine), PHE (phenylalanine), TYR (tyrosine), HIS (histidine), DHAP (dihydroxyacetonephosphate), S7P (sedoheptulose-7-phosphate) plotted on a metabolic map showing reactions for glycolysis, PPP, GLX (glyoxlyate shunt) and the TCA cycle. Aspartate/asparagine and glutamate/glutamine pools are lumped as both asparagine and glutamine were reduced to aspartate and glutamate respectively during acid hydrolysis.

For steady state CHL-ACE Mtb cultures, the ^13^C labelling profiles reflect the entry points of cholesterol into central carbon metabolism. Cholesterol catabolism yields four propionyl-CoA, four acetyl-CoA, one pyruvate, and one succinyl-CoA of which succinyl CoA, pyruvate and acetyl-CoA enters central carbon metabolism directly (28). Propionyl-CoA is toxic to Mtb and can be metabolised by either: (1) the methyl citrate cycle to pyruvate (2) the B12 dependent methyl malonyl pathway leading to succinyl-CoA or (3) used in cell wall lipogenesis. In our experiments the methyl malonyl pathway is not active (as Roisin’s media lacks vitamin B12).

For the CHL-ACE experiments, the labelling profile of succinate (SUC), pyruvate (PYR), alanine (ALA), valine (VAL) and leucine (LEU) had ≥ 50% unlabelled carbon indicating that the carbon backbone of these metabolites was predominantly derived from unlabelled cholesterol. Canonical ^13^C labelling patterns were measured in the other metabolites reflecting that 60% of the total carbon was derived from ^13^C labelled acetate and the remainder derived from unlabelled cholesterol indicating that metabolism was also not compartmentalised in these conditions (Fig. 1). For GLY-OLA grown cultures, the labelling profiles was consistent across the different metabolites analysed; the backbone of these metabolites was synthesized primarily from glycerol, demonstrating that metabolism of GLY-OLA was also not compartmentalised.

### ^13^C isotopologue analysis of amino acids provides their distinct biosynthetic routes

We performed ^13^C isotopologue analysis of proteinogenic amino acids. As expected the profiles of all metabolites derived from CHL-ACE were different to that derived from GLY-OLA grown Mtb (Fig. 2). The labelling profiles of amino acids reflect their biosynthetic origin for both GLY-OLA and CHL-ACE grown Mtb. For the GLY-OLA cultures, the labelling patterns of aspartate (ASP/N), threonine (THR), isoleucine (ISO), lysine (LYS) and methionine (MET) were similar, consistent with a common biosynthetic origin of these amino acids from oxaloacetate. The isotopologue composition of tyrosine (TYR) and phenylalanine (PHE) were also alike reflecting their common precursors-phosphoenolpyruvate and erythrose-4-phosphate. Similarly, ornithine (ORN) and glutamate (GLU/N), which are both derived from α-ketoglutarate had similar labelling patterns. The isotopologue profiles of proteinogenic amino acids derived from steady state cultures grown in CHL-ACE also reflect their biosynthetic origin. THR and MET had near identical profiles with ASP/N with M+2 and M+4 isotopomers having the highest ^13^C labelling, indicating their synthesis from ASP. GLU/N and ORN, TYR and PHE and ALA and SER all had very similar profiles indicating the common biosynthetic precursor for each of these pairs of amino acids. Whilst manual analysis of ^13^C isotopologue data allows qualitative conclusions, systems level analysis is required in order to quantitate the metabolic fluxes.

**Fig. 2.**
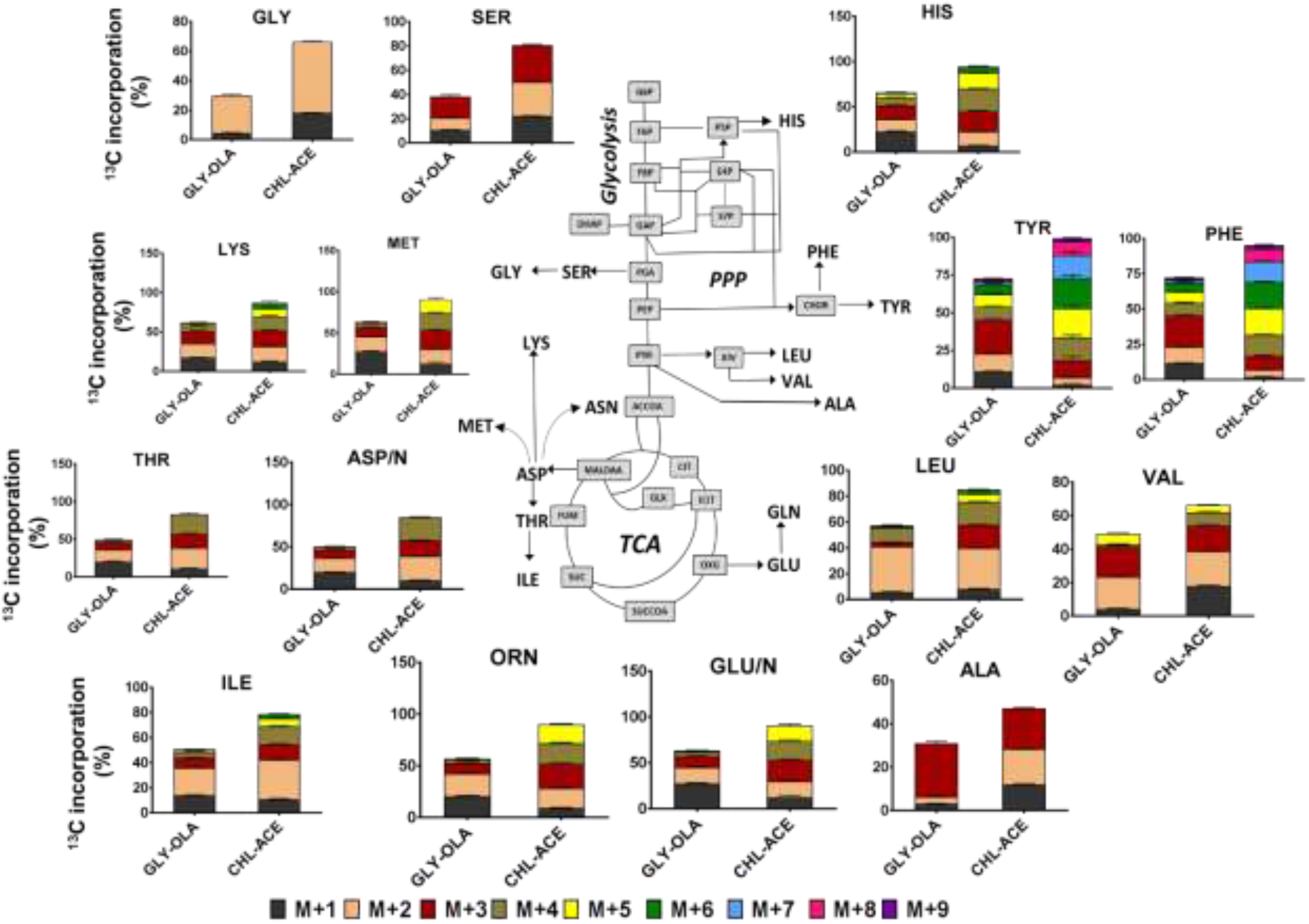
^13^C isotopologue analysis of proteinogenic amino acids from Mtb grown on CHL-ACE and GLY-OLA. Measurements were obtained from Mtb grown under steady state chemostat conditions. ^13^C incorporation is shown for the mass isotopomers, which are labelled as ^13^C_1_, ^13^C_2_ etc. Measurements are shown for alanine (ALA), leucine (LEU), valine (VAL), glycine (GLY), serine (SER), methionine (MET), histidine (HIS), aspartate-asparagine (ASP/N), threonine (THR), lysine (LYS), isoleucine (ILE), glutamate-glutamine (GLU/N), ornithine (ORN), tyrosine (TYR) and phenylalanine (PHE. Glycolysis, pentose phosphate pathway (PPP), glyoxylate shunt (GLX) and the TCA cycle are depicted. Values are mean ± S.D. of three replicates.

### *In vivo* flux profiles of Mtb obtained with ^13^C-metabolic flux analysis

^13^C-MFA is the preferred tool for quantitative characterization of metabolic phenotypes in steady-state cultures (4, 27). Previously we developed an isotopomer model of Mtb’s central carbon metabolism comprising of the TCA cycle, glycolysis, pentose phosphate pathway (PPP) and anaplerotic reactions, which allowed us to analyse ^13^C-isotopomer data and compute metabolic fluxes (4). For this study we have expanded our capacity for predicting metabolic fluxes by adding the methyl citrate cycle (MCC) and amino acid degradation pathways to our previous ^13^C-isotopomer model (see Table S1 in the supplementary material for network details). Absolute flux distributions were derived from the two steady state chemostat conditions (Fig. 3) using the INCA platform (28). In our previous study we were unable to determine the flux values unambiguously for Mtb growing in GLY-OLA. Here by using our extended isotopomer model we obtained a unique metabolic flux profile for these conditions building on our previous work (Fig. 3). A comparison of the current and the previous metabolic flux profiles of Mtb growing in GLY-OLA in steady state cultures (Fig. S2) illustrates the similarities between the solutions and shows that extending the isotopomer model has improved our ability to resolve the metabolic fluxes of Mtb.

**Fig 3.**
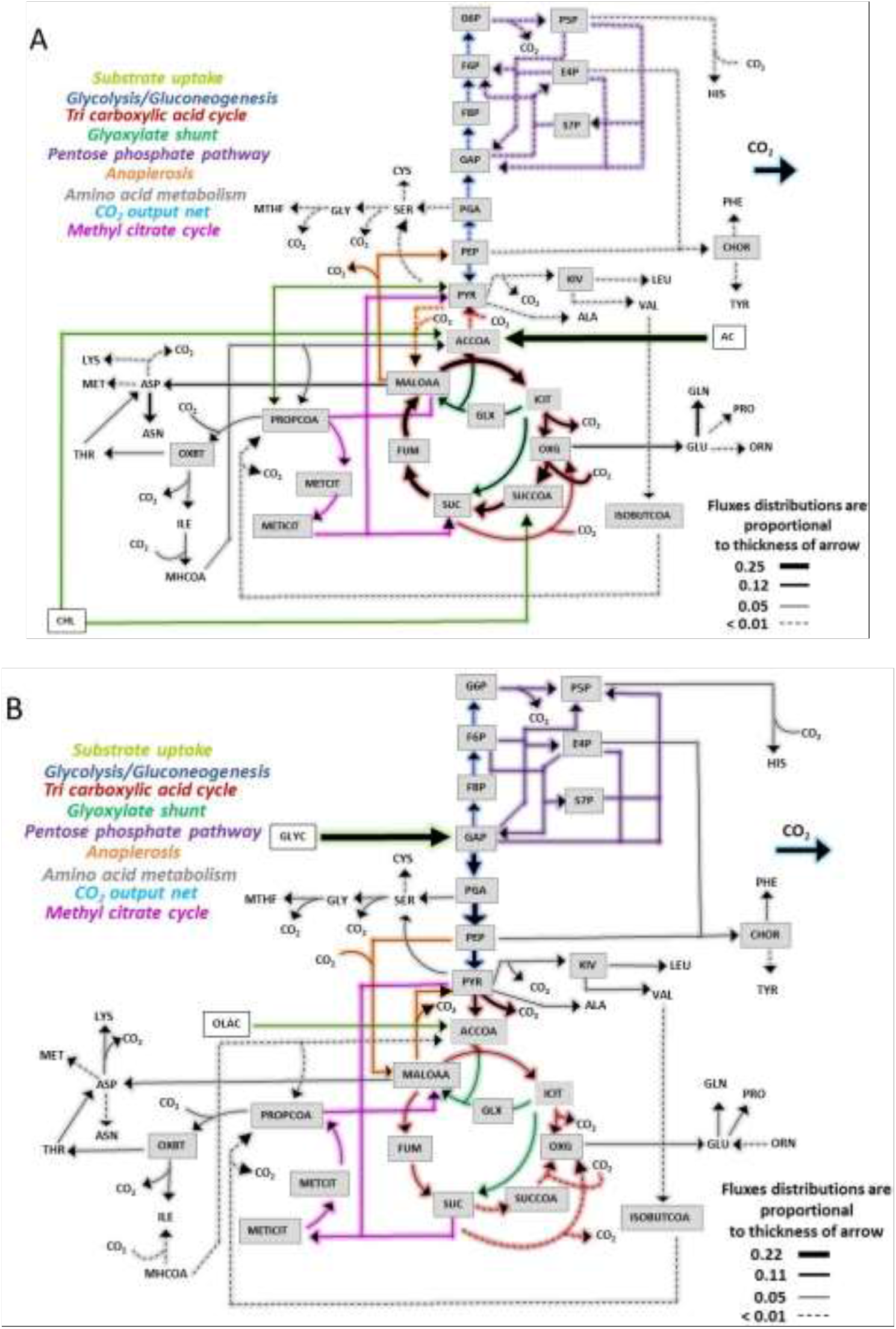
Flux distributions of Mtb grown on CHL-ACE (A) and GLY-OLA (B) carbon substrate combinations. Flux distributions were calculated for the extended Mtb metabolic network shown in supplementary Table S1. Fluxes are absolute values expressed as mmol g^-1^ biomass h^-1^. ^13^C labelled substrate consumption-acetate (ACE) and glycerol (GLYC) and CO_2_ production fluxes were fixed for the calculations. Steady state ^13^C mass isotopomer measurements for the protein-derived amino acids from CHL-ACE and GLY-OLA grown Mtb were used for flux estimations. Estimated fluxes in supplementary data file S2 are represented by arrows in Figure 3A and B. The width of each line is proportional to the underlying flux value. Metabolite lists for the flux maps are included in supplementary data file S2.

### Distinct metabolic flux partitioning between the TCA cycle and glyoxylate shunt during growth on cholesterol-acetate and glycerol-oleic acid

The resolved flux distributions of Mtb growing on CHL-ACE demonstrated that there is a significant difference between how Mtb partitions carbon flux between the TCA cycle and the glyoxylate shunt in CHL-ACE as compared with GLY-OLA (Fig. 3A). When growing on CHL-ACE Mtb operates a complete oxidative TCA cycle in combination with an active glyoxylate shunt. There is only a very low flux through pyruvate dehydrogenase (PDH) (TCA R1) with the small amount of cholesterol derived pyruvate being channelled to leucine, valine and alanine biosynthesis (Fig. 3A, 4A) In concordance with our previous work, Mtb uses an incomplete TCA cycle when growing on GLY-OLA (4).

**Fig. 4.**
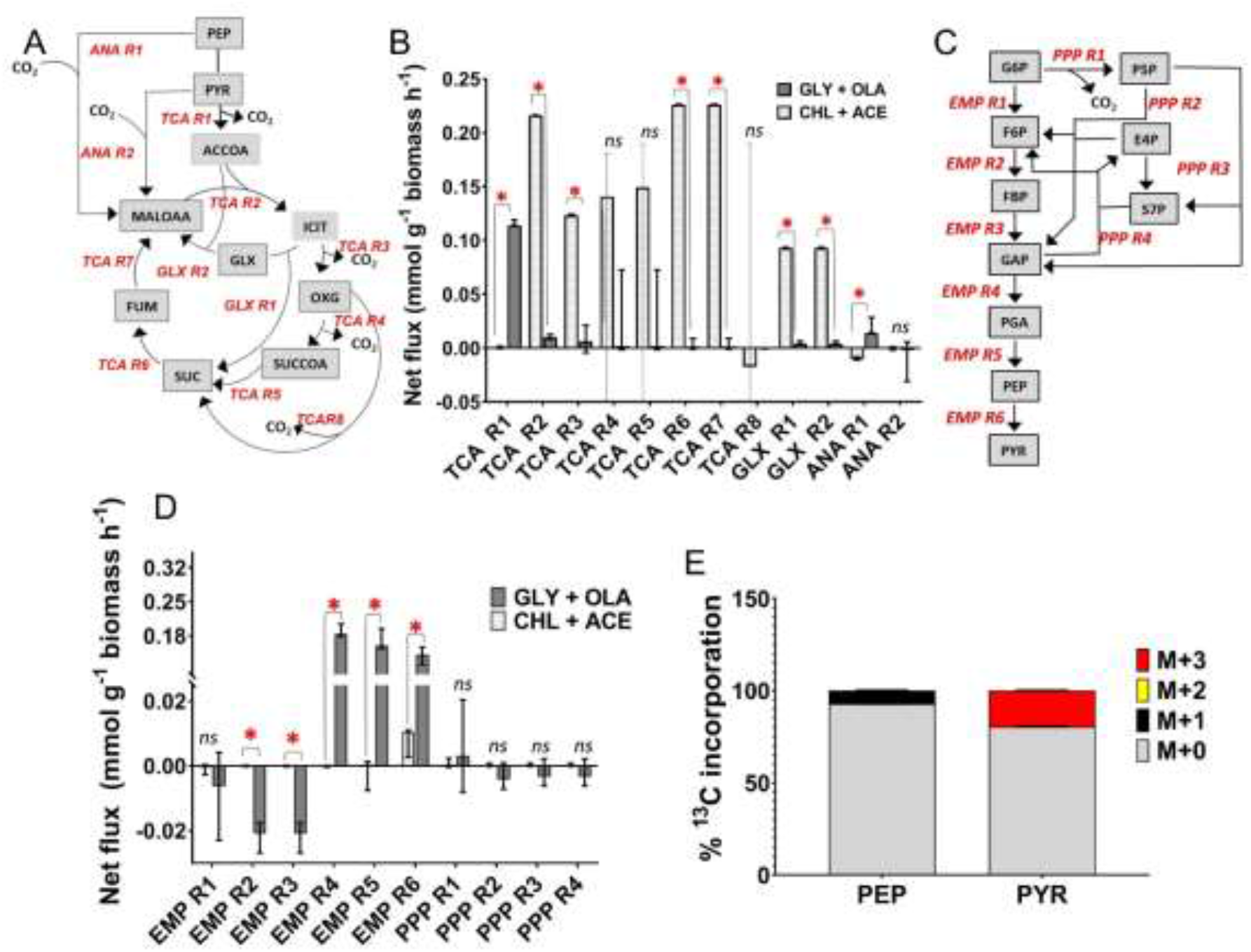
Flux distributions through the TCA cycle, glyoxylate shunt, anaplerotic pathways, glycolysis and gluconeogenesis for Mtb grown on CHL-ACE and GLY-OLA. The network reactions for the TCA cycle (TCA R1, TCA R2, TCA R3, TCA R4, TCA R5, TCA R6, TCA R7), glyoxylate shunt (GLX R1, GLX R2) and anaplerotic reactions (ANA R1, ANA R2) along with their directionalities are shown in (A). (B) Quantitative comparisons of the best-fit net fluxes.The error bars on these measurements show the lower and upper confidence limits. (C) The network reactions for glycolysis (EMP R1, EMP R2, EMP R3, EMP R4, EMP R5, EMP R6) and for PPP (PPP R1, PPP R2, PPP R3, PPP R4) along with their directionalities.(D) The best-fit net fluxes for glycolysis and PPP respectively. Statistically significant fluxes are indicated by *;ns denotes not significant. (E) ^13^C incorporation in pyruvate (PYR) and PEP of Mtb grown in [3, 4-^13^C_2_] cholesterol. A negative flux indicated the reverse directionality of the flux to that shown in Figures A and D. Metabolites shown in A and D are MALOAA (malate + oxaloacetate), SUC (succinate), ACCOA (acetyl coenzyme A), PYR (pyruvate), ICIT (isocitrate), GLX (glyoxylate), OXG (α-ketoglutarate), SUCCOA (succinyl coenzymeA), FUM (fumarate), G6P (glucose-6-phosphate), F6P (fructose-6-phosphate), FBP (fructose 1,6-bisphosphate), GAP (glyceraldehyde-3-phosphate), PGA (phosphoglyceric acid), PEP (phosphoenolpyruvate), PYR (pyruvate), P5P (pentose-5-phosphate), E4P (erythrose-4-phosphate), S7P (sedoheptulose-7-phosphate).

Determining the individual anaplerotic fluxes from ^13^C-isotopomer data presents a major challenge. For example, it is not possible to discriminate between the reactions of malic enzyme (MEZ) and pyruvate carboxylase (PCA) with high confidence so the summed flux is shown between pyruvate and malate/oxaloacetate (ANA R2, Fig 4). From the data we identified that Mtb uses phosphoenolpyruvate carboxykinase (PEPCK or ANA R1) gluconeogenically during growth on CHL-ACE to generate PEP, which is in accordance with the use of the glyoxylate shunt for replenishing the TCA intermediates. Whereas, when growing on GLY-OLA, PEPCK operates in the carboxylating direction to fix CO2 as we have described previously (4). The overall flux through ANA R2 is very low and not significantly different when growing on CHL-ACE or GLY-OLA.

Perhaps, surprisingly, there was no flux from pyruvate to PEP, the reaction catalysed by the fourth member of the anaplerotic node, pyruvate phosphate dikinase (PPDK), in either condition. There was flux towards pyruvate mediated by pyruvate kinase (PYK = EMP R6) and this was higher on GLY/OAA than with CHL/ACE (Fig. 4C, D). We previously demonstrated using mutagenesis that Rv1127c (annotated as PPDK) was essential for the growth of Mtb in rich media containing cholesterol even in the presence of an additional growth permissive carbon source (including acetate). This phenotype was associated with the inability of the knockout Mtb strain to metabolise the cholesterol by-product propionate, which is toxic to bacterial cells (23). High through-put Tn screen also identified Rv1127c as required for growth on cholesterol. However, other studies suggest that Mtb lacks PPDK activity (21). In order to explore this further we performed a ^13^C labelling experiment growing Mtb *in vitro* on Roisin’s minimal medium containing [3, 4-^13^C_2_] cholesterol as a sole carbon source (Fig. 4E) This experiment identified [^13^C_2_] pyruvate, as expected when cholesterol is catabolised, however all of the PEP detected was only labelled in one carbon (M+1), indicating that the other ^13^C label has been lost as CO2 via the TCA cycle consistent with the metabolic flux prediction that Mtb lacks PPDK activity (Fig. 4D, E).

We observed only a very low gluconeogenic flux (Fig. 4C and D) during growth on CHL-ACE, which was in sharp contrast to the glycolytic fluxes from glycerol to pyruvate, when grown on GLY-OLA. Pyruvate kinase (PYK) was however active under both conditions, where it funnels the PEP generated from PEPCK to pyruvate when growing on CHL-ACE (Fig. 4C, D). On GLY-OLA the flux through PYK was higher, a prediction which agrees with reports that this enzyme is required for carbon co-metabolism (29). Fluxes through the pentose phosphate pathway (PPP) were much lower when Mtb was grown with CHL-ACE as compared to GLY-OLA. The PPP not only supplies pentose phosphates, but also reducing power in the form of NADPH which can then be used for lipid biosynthesis and other anabolic processes. This may reflect a higher demand for NADPH from the PPP when growing Mtb is growing on GLY-OLA as compared to cultures containing the highly reduced carbon source, cholesterol (Fig. 4C, D).

### Methyl citrate cycle fluxes are reversed during metabolism of glycerol-oleic acid but not on cholesterol-acetate

The MCC in mycobacteria is required for the utilization of the propionyl-CoA derived from sterols, branched chain amino and fatty acids that are acquired from the host (Fig. 5A). Here we show that when Mtb is growing on cholesterol and acetate there is very low flux through the MCC suggesting that this pathway is minimally required for cholesterol metabolism in the presence of acetate (Figure 5A, B). Previous studies have demonstrated that the necessity for the MCC can be overridden by supplying Mtb with AcCoA generating carbon sources which primes the conversion of propionyl-CoA into methyl malonyl-CoA-lipids such as sulfolipid-1 (SL-1), polyacyltrehalose (PAT), and phthiocerol dimycocerosate (PDIM) (25). This lipid synthesis may account for the increased extracellular biomass observed in CHL/ACE grown cells (Table 1). By contrast when Mtb is growing on GLY-OLA, there is a reversal of flux through the MCC (MCC R1-R3) presumably to generate propionyl-CoA for lipid biosynthesis, which is also supplemented by the degradation of the branched chain amino acid valine (D R1 and D R2) (Fig. 5B-D).

**Fig 5.**
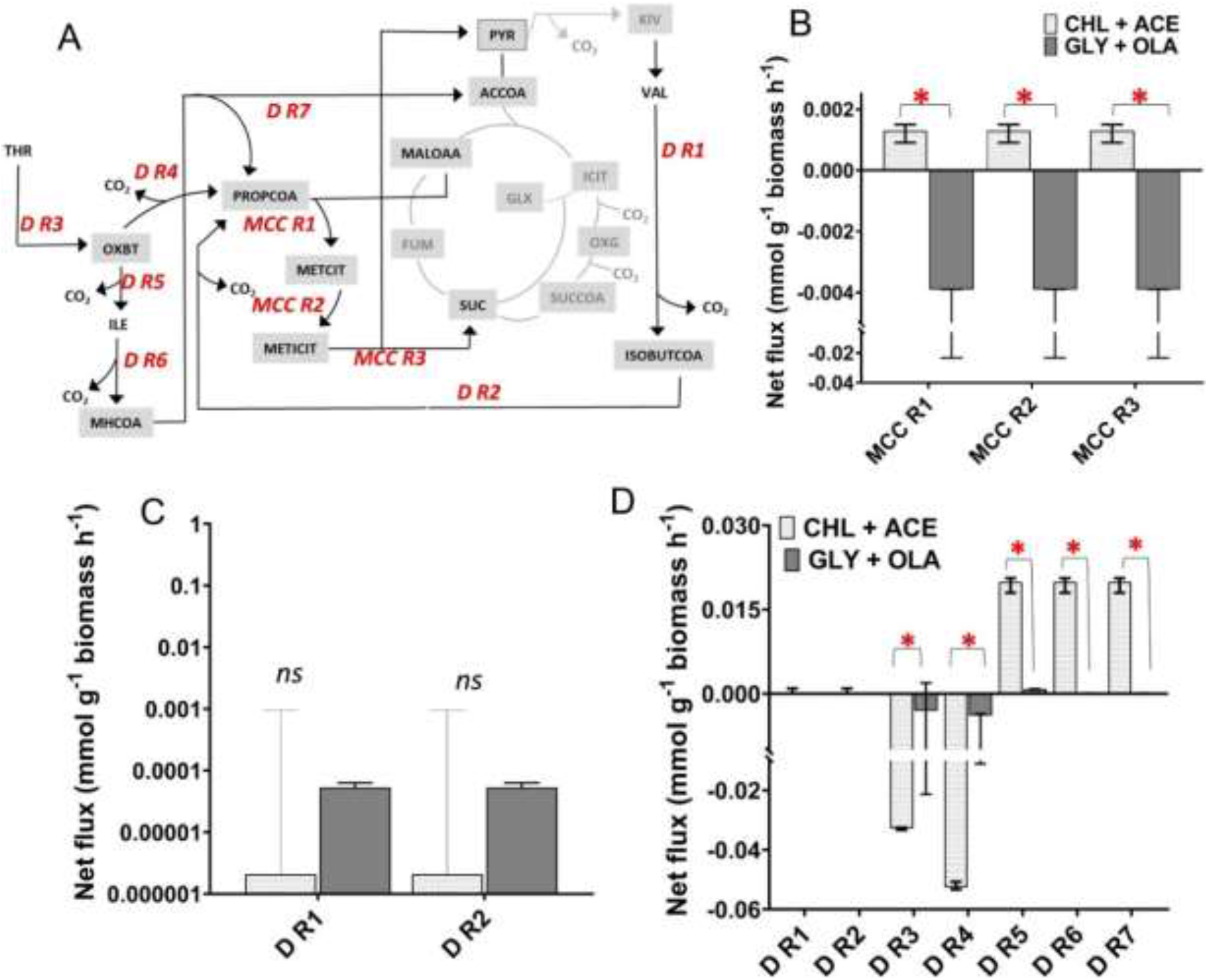
Flux distributions through the methyl citrate cycle and amino acid degradation pathways of Mtb grown on CHL-ACE and GLY-OLA. (A) The network reactions for methyl citrate cycle (MCC R1, MCC R2, MCC R3), amino acid degradation pathways (D R1, D R2, D R3, D R4, D R5, D R6, D R7) along with their directionalities are shown in A. (B) Best-fit absolute net fluxes of the methyl citrate cycle and the error bars are lower and upper confidence limits of the net fluxes (C) Best-fit absolute net fluxes for valine degradation, threonine and isoleucine degradation on CHL-ACE and GLY-OLA. A negative flux indicated the reverse directionality of the flux to that shown in A. The error bars on these measurements show the lower and upper confidence limits for the fluxes. Statistically significant differences for the fluxes measured on the two growth substrates are indicated by *; ns denotes not significant. A negative flux indicated the reverse directionality of the flux to that shown in A. Abbreviations for metabolites are MALOAA (malate + oxaloacetate), SUC (succinate), ACCOA (acetyl coenzyme A), PYR (pyruvate), METCIT (methyl citrate), METICIT (methyl isocitrate), MMCOA (methyl malonyl coenzyme A), ISOBUTCOA (isobutyl coenzyme A), OXBT (oxobutanoate), ILE (isoleucine), THR (threonine), MHCOA (methylacetoacetyl coenzyme A).

In order to explore the incorporation of carbon into lipids, we performed MALDI-TOF analysis on Mtb from our steady state chemostat cultures (Fig. 6). This data cannot be incorporated into flux analysis as there is no one-to-one mapping between carbon atoms in lipids and precursor metabolites. We identified a large number of complex lipid species, including phosphatidylinositol mannosides (PIMs) and SL-1 (Fig.6) In order to test whether this was due to incorporation of ^13^C acetate into SL-1 or an increased chain length due to the incorporation of carbon from unlabelled cholesterol, as predicted by our flux profiles, we also measured the lipid fingerprint of control samples derived from the same chemostat cultures but prior to attaining an isotopic steady state. In accordance with our metabolic flux predictions we identified an increase in chain length in SL-1 in these unlabelled control samples which confirmed our expectations that lipids derived from CHL-ACE-grown Mtb had significant differences in the composition of the methyl malonyl-CoA derived SL-1s, specifically an increase in chain length and abundance when compared to GLY-OLA grown Mtb (Fig.6) The increase in chain length in CHL-ACE grown Mtb indicates incorporation of carbons into the SL-1s from cholesterol that has a higher number of carbon atoms than glycerol (27 carbons). This is concordance with other studies which show an increased chain length of SL-1 during growth of Mtb on cholesterol and propionate (24). We also found significant mass shifts in triacylphosphatidylinositol dimannosides (AcPIMs) isolated from the chemostat samples. However these shifts were not observed in the control samples indicating that the in this case the increased mass was due to ^13^C labelled acetate or glycerol being incorporated into these lipids. For CHO-ACE-grown Mtb, the observed increased mass shift (Fig. S3A, B) was likely due to incorporation of [^13^C_2_] acetate) directly into these lipids in contrast to the GLY-OLA chemostat grown Mtb where unlabelled OLA was being predominantly used for acetylation (Fig. S3).

**Fig 6:**
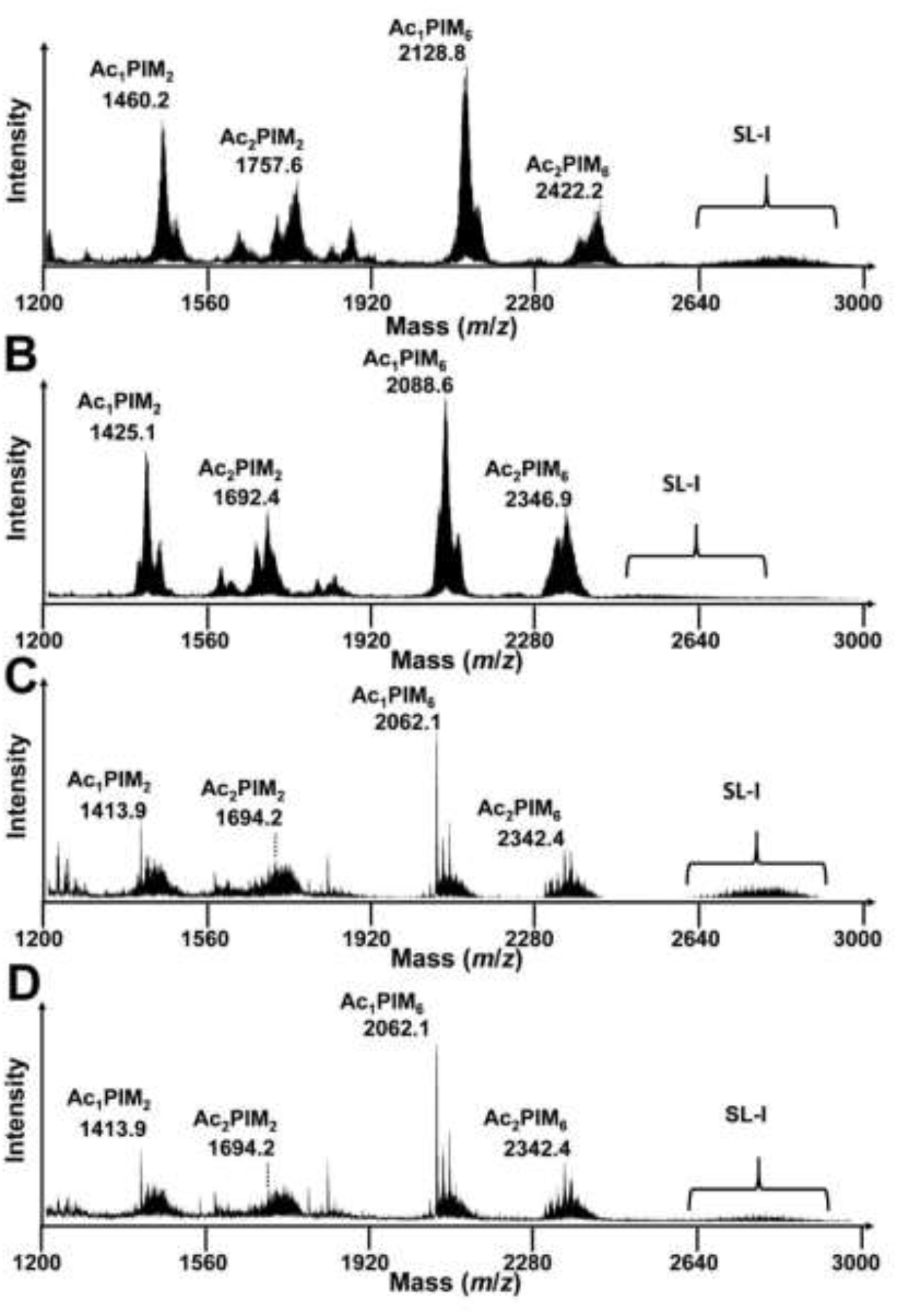
Negative ion mode MALDI-ToF mass spectra showing the lipid fingerprint of Mtb under different carbon sources in chemostat. Ac1PIM2: tri-acyl phosphatidylinositol mannoside di-mannose; Ac2PIM2: tetra-acyl phosphatidylinositol mannoside di-mannose; Ac1PIM6: tri-acylated phosphatidyl-myo-inositol hexamannoside; Ac2PIM6: tetra-acylated phosphatidyl-myo-inositol hexamannoside and SL-I: sulpholipid. The ion at m/z 1413.9 is assigned to the deprotonated PIM2 containing 2 C16:0 and 1 C19:0; The ion at m/z 1694.1 is assigned to the deprotonated PIM2 containing 2 C16:0 and 2 C19:0; The ion at m/z 2062.1 is assigned to the deprotonated PIM6 containing 2 C16:0 and 1 C19:0; The ion at m/z 2342.9 is assigned to the deprotonated PIM2 containing 2 C16:0 and 2 C19:0; SL-I is observed as a broad set of peaks representing multiple lipoforms corresponding to differing numbers of CH_2_ units. Multiple lipoforms of SL-1 (m/z 2500 to m/z 3000) are also indicated SL-I. Mass shifts due to ^13^C labelling of whole bacteria lipid fingerprint prepared from metabolic and isotopic steady state chemostat Mtb grown with [^13^C_2_] acetate and unlabelled cholesterol (A), [^13^C_3_] glycerol and unlabelled Tween80 (B), unlabelled acetate and (C) and unlabelled glycerol and Twen80 (D). Samples taken from the same chemostat at metabolic steady state (A) and (B) and isotopic in stationary state (C) and (D). The resolved mass shifts observed for Ac1PIM2 are shown to indicate significant lipid changes between the two conditions.

### Mtb uses cholesterol to synthesize amino acids

Since the synthesis of amino acids is required for biomass production we also calculated the amino acid production rates. ^13^C-MFA demonstrated that there were significant differences in the amino acid biosynthesis fluxes between the two chemostat conditions. For example the synthesis of ASP/ASN, GLU/GLN and ILE were significantly higher for Mtb growing slowly with CHL-ACE than when the carbon sources were GLY-OLA (Fig. 7A). Scrutinising this data further shows that the fluxes though GLU/GLN synthesis ultimately contribute towards asparagine biosynthesis. This is in concordance with the data showing that asparagine accumulates during metabolism of the cholesterol by-product propionate (30).

**Fig. 7:**
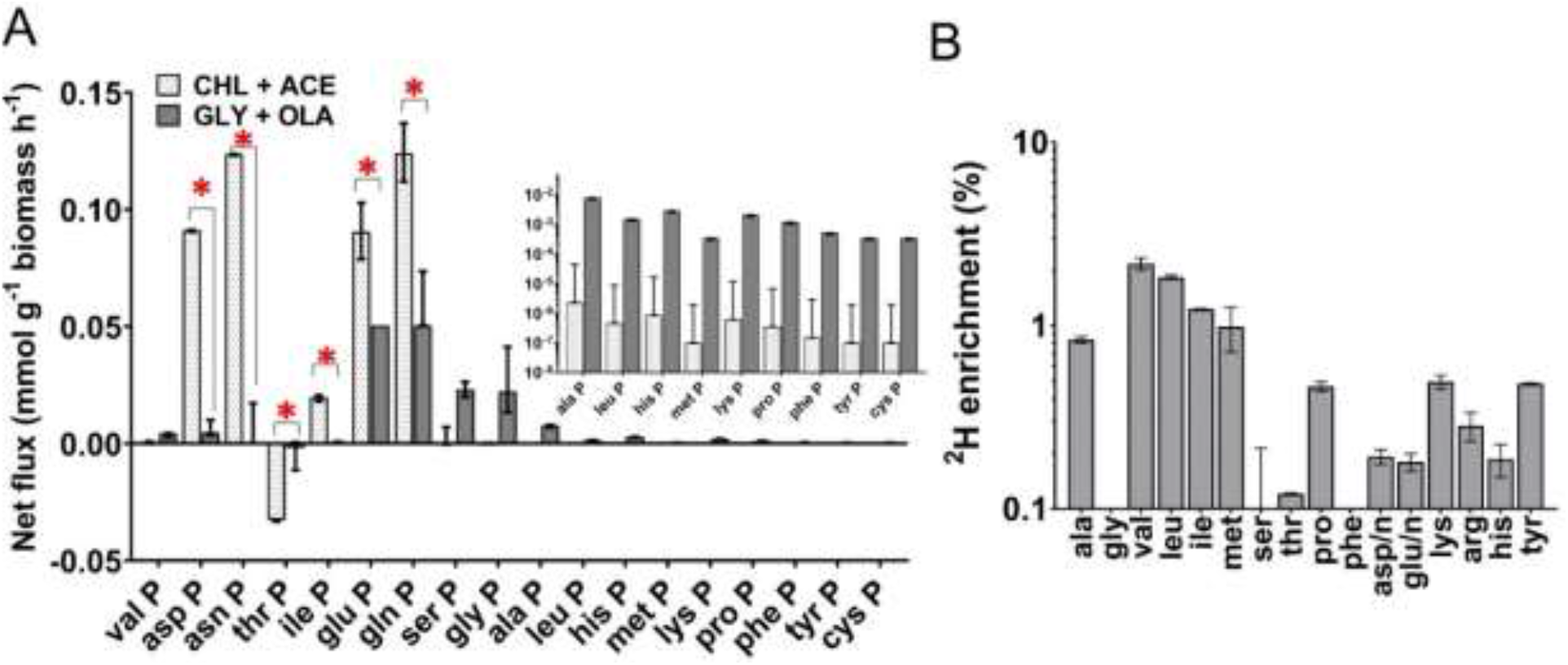
Amino acid biosynthesis. A) Fluxes are shown for the synthesis of amino acids alanine (ala P), valine (val P), leucine (leu P), tyrosine (tyr P), phenylalanine (phe P), histidine (his P), glutamine (gln P), glutamate (glu P), proline (pro P), ornithine (orn P), asparagine (asn P), aspartate (asp P), threonine (thr P), isoleucine (ile P), methionine (met P), lysine (lys P), glycine (gly P), cysteine (cys P) and serine (ser P). Statistically significant differences for the fluxes measured on the two growth substrates are indicated by * **B)** ^2^H (deuterated hydrogen) incorporation in the amino acids of Mtb grown in minimal media with deuterated cholesterol as the sole carbon source. Values are mean ± S.E.M (n=3 biological replicates).

The increase in biosynthesis of ILE during growth on CHL-ACE is of interest as in other fungi and bacteria it has been shown that branched chain amino acid biosynthesis can also function as a mechanism to dissipate reductants (31–32) Supported by the lipid profiling data here it is thought that Mtb primarily uses lipids as an electron sink to maintain redox homeostasis when growing on highly reduced carbon sources such as cholesterol (24–26). However the role of amino acids as potential electron sinks has never been explored for Mtb. Thus we grew Mtb in Roisin’s minimal medium containing uniformly deuterated cholesterol and measured the incorporation of deuterium into amino acids. This analysis showed that Mtb incorporates hydrogen from cholesterol into several amino acids with branched chain amino acids having the highest level of incorporation (Fig. 7B). This data suggests that amino acid anabolism could provide Mtb with another mechanism for sinking electrons and maintaining redox homeostasis when consuming highly reduced carbon sources such as cholesterol.

## Discussion

Mtb is known to be able to acquire and assimilate multiple carbon sources from the host during infection (5, 6, 13, 19). This co-catabolism has been demonstrated to occur both *in vitro* and during intracellular growth (14). Details of the metabolic mechanisms involved in this process have been inferred from ^13^C labelling experiments in which specific nutrients were observed to have defined fates suggesting some form of compartmentalization of metabolites (14). However, in our steady state, slow growing cultures we demonstrated efficient co-catabolism of either cholesterol and acetate, or glycerol and oleic acid but found no evidence of compartmentalised metabolism. This apparent discrepancy suggests that metabolic compartmentalisation may be carbon source-dependent, or may reflect differences in experimental setup. The previous studies were performed using Mtb grown in non-metabolic steady state conditions on Middlebrook 7H10 agar containing in addition to the carbon sources (one of which was ^13^C labelled) significant amounts of unlabelled glutamate which Mtb can use as both a nitrogen and a carbon source (33). The instationary metabolic state of the culture would affect the rates of ^13^C incorporation into intermediates whilst the additional unlabelled carbon source would have diluted out the label thus potentially confounding the analysis.

Compartmentalisation in prokaryotic cells can involve micro-compartments or cages. These are semi-permeable protein shells that sequesters enzyme complexes from a segment of a metabolic pathway, but that are permeable to metabolites. These structures allow pathway reactions to be spatially organised and so can enhance flux through multi-step pathways, or protect cells from toxic metabolic intermediates (34–35). To date, only relatively small cages have been identified in mycobacteria, which encapsulate pathways involved in oxidative stress and iron storage pathways (36). Mtb does not encode the various subunits required to make-up the larger compartments that could, for example, encage whole metabolic pathways. Therefore, with our current knowledge, it is difficult to understand how Mtb could segregate metabolites from single carbon sources through a large pathway such as the TCA cycle for example. The hypothesis that Mtb compartmentalises central carbon metabolism thereby requires further investigation.

The physiological role of Mtb’s glyoxylate shunt is to supply and replenish the TCA and MCC cycle intermediates, oxaloacetate, to ensure adequate supply for gluconeogenesis and amino acid synthesis by bypassing the decarboxylating steps of the TCA cycle and thereby preserving carbon (37, 38). The pathway is now known to have a multifaceted role in microbial growth and survival in a variety of conditions and is also critical for the virulence of a number of pathogens including Mtb (4, 26, 39). During growth on CHL-ACE, Mtb had significantly higher fluxes through both the complete TCA cycle and the glyoxylate shunt than when grown with GLY-OLA.

In the highly oxygenated chemostat, cholesterol catabolism will generate reductants that need to be recycled (35–37). Our flux results demonstrate that during slow growth with CHL-ACE this is occurring via the TCA cycle and then through oxidation of NADH in the electron transport chain (28–29, 40). Our data also indicates that lipids provide a sink for reductants in these conditions as has been described previously (26, 41). However, our studies using deuterated cholesterol indicate that some reductant is also channelled into amino acid biosynthesis indicating that this provides Mtb with another potential sink for reductants.

The relatively large amounts of CO2 produced from the CHL/ACE culture is in concordance a partitioning of isocitrate through a complete TCA cycle and the glyoxylate shunt combined with gluconeogenic operation of PEPCK when growing on cholesterol and acetate. This mode of operation is in stark contrast to Mtb growing with GLY-OLA where we measured reduced flux through an incomplete TCA cycle and insignificant fluxes through the glyoxylate shunt combined with anaplerotic flux through PEPCK that leads to the lower CO2 release in Mtb grown on GLY-OLA. OAA supply is in part satisfied by PEPCK functioning in the carboxylating direction, confirming our previous result (4), whilst gluconeogenesis proceeds directly from glycerol derived PGA. Although MEZ, PCA, (which are not distinguished in our model) and PEPCK are all able to fix carbon from CO2 (4, 42). PEPCK is predicted by these results to be the dominant enzyme of the anaplerotic node required for either gluconeogenesis or anaplerosis. This explains why Mtb strains lacking PEPCK are attenuated in a variety of different *in vitro, ex vivo* and *in vivo* conditions (5, 29, 43).

Previous studies have shown that Mtb strains lacking the bifunctional isocitrate lyase/methylisocitrate lyase (ICL) are unable to grow in the presence of cholesterol because they cannot metabolise propionate due to the role of this enzyme in the MCC (25). However, the addition of C2 containing metabolites alleviates this attenuation, by priming incorporation of propionyl-CoA into lipids and so limiting the build-up of toxic MCC intermediates and propionyl-CoA itself (25). Here we show that even in the presence of an intact MCC the flux of carbon from cholesterol into lipids is the preferred metabolic route when there are sufficient C2 metabolites to prime lipid biosynthesis as evidenced by the low flux though the MCC in the slow growth chemostat conditions. This was further confirmed by the lipid analysis which showed an increase in the incorporation of carbon from cholesterol into SL-1 resulting in an increase in chain length of this virulence lipid.

The role of the glyoxylate shunt in Mtb pathogenesis has always been obscured by the duel function of the gating enzyme ICL enzyme in both the shunt and the MCC. Enzymes which function exclusively in the MCC are known to be dispensable in murine models of Mtb, prompting the hypothesis that vitamin B12 dependent methyl malonyl pathway (MMP) complements for the absence of the MCC pathway *in vivo* (12, 23). However this is not consistent with the finding that strains lacking both a functional MCC and MMP but with a functioning glyoxylate shunt were significantly less attenuated in a murine model of TB than Δ*icl* Mtb strains (11). This indicates that in some conditions ICL’s only role is as the gateway enzyme into the glyoxylate shunt. Here we show that when Mtb is growing slowly with cholesterol and fatty acids there is very low flux through the MCC and the main function of ICL is to channel carbon through the glyoxylate shunt.

We previously identified an additional essential role for ICL during slow growth of Mtb on glycerol and Tween80 (4) using ^13^C MFA, conditions in which the glyoxylate shunt would not be expected to be active (4). Here, using our extended isotopologue model, we demonstrate that under these conditions ICL is catalysing a reverse flux through the MCC in the direction leading to propionyl-CoA generation. Reversal of the MCC has been described by Serafinini *et al*. when Mtb is growing with pyruvate or lactate (44). Our data suggests that this is a more general phenomena and may be active whenever there is a limited supply of odd-chain lipid precursors. The ability to operate the MCC as both a catabolic and anabolic pathway provides Mtb with the metabolic flexibility to both detoxify excess sterol-derived propionate, and also, when necessary, to generate propionyl-CoA for lipid biosynthesis, including virulence-associated lipids. The generation of propionyl-CoA maybe may be particularly important when Mtb is growing slowly and in stress conditions which are associated with increased demand for specific cell wall lipids (24, 45–47). This also suggests that in conditions where a reversed MCC is required the methyl malonyl pathway would be unable to complement the MCC pathway, highlighting the necessity for both pathways.

A role for a reversed MCC in the co-metabolism of glycerol and fatty acids is of interest in the context of the discovery that Mtb has access to glycerol *in vivo* (19). Although unlikely to be an essential carbon source, changes in glycerol metabolism have been shown to influence antibiotic tolerance of Mtb *in vivo* (17, 48). Also, mutations in the MCC regulator *prp*R that conferred multidrug tolerance were identified in TB patients (49). Although this finding was proposed to reflect an impaired ability of these strains to metabolise propionate, the results here suggest an alternative and intriguing hypothesis that dysregulation of the MCC may affect the metabolism of glycerol. Interestingly, Mtb strains lacking ICL were shown to be less tolerant to antibiotics in glycerol containing medium and this was associated with an increased flux through the TCA cycle (21, 50). We posit that this is due to the operation of a reversed MCC. When the MCC is reversed methyl citrate lyase (ICL) becomes the gating enzyme into this cycle and therefore as Nandakumar *et al* eloquently demonstrated ΔiclMtb strains would increase flux through the TCA cycle, altering the energetics of the cell and reducing Mtb tolerance to antibiotics (50).

Surprisingly, our data predicted zero flux through the PPDK reaction presumed to be catalysed by the product of Rv1127c and has been shown by us, and others, to be essential for cholesterol and propionate metabolism (21, 42). PPDK’s use two binding domains, for ATP/AMP and PEP/pyruvate, to catalyse the reversible interconversion between pyruvate and PEP. Mysteriously, analysis of Rv1127c reveals that although the gene encodes an entire N-terminal PPDK domain to bind and dephosporylate ATP, the predicted protein completely lacks a C-terminal PEP/pyruvate binding domain (Fig. S4), raising doubts over the annotated function of this protein. *Mycobacterium bovis, Mycobacterium bovis* BCG and *Mycobacterium leprae* encode similar “PPDK” genes lacking PEP/pyruvate binding domains (33, 51–52). Although it is possible that the mycobacterial PEP/pyruvate binding domain is encoded on another Mtb gene, *in silico* searches have not identified any genes Mtb protein homologous to the C-terminal PEP/pyruvate-binding domain of typical PPDK’s. Loci for functionally related genes tend to be in close genomic proximity in prokaryotes and the Rv1127c locus is nearby genes encoding enzymes and regulators of the methylcitrate cycle that, like Rv1127c, are essential for the metabolism of cholesterol and propionate. The absence of PEP/pyruvate binding domain and, as we demonstrate in this study, zero flux through the PPDK reaction in conditions in which Rv1127c is essential, prompts us to propose that Rv1127c is not a PPDK but performs an, as yet unknown, metabolic function associated with the methylcitrate cycle. This would be consistent with our previous data demonstrating propionate vulnerability of strains lacking Rv1127c that could be complemented by vitamin B12 to activate the methyl-malonyl pathway. The precise function of Rv1127c may have therapeutic implications as ΔRv1127c is attenuated for intracellular survival and is more sensitive to the antibiotic bedaquiline (BDQ) (53). Rv1127c is thereby as an attractive target for developing anti-TB drugs which synergise with bedaquiline, a critical drug in the new shortened therapy against drug resistant TB.

In conclusion our results demonstrate that Mtb efficiently co-metabolises combinations of either glycerol or cholesterol along with C2 generating substrates to efficiently generate biomass. Although it has been suggested that Mtb is able to differentially co-metabolise carbon substrates to distinct metabolic fates we find no evidence of compartmentalised metabolism in this study suggesting that this hypothesis may need re-evaluating. We show that a reversible MCC pathway along with the glyoxylate shunt provides Mtb with a high level of flexibility for re-routing metabolic fluxes depending on the combination of carbon sources being metabolised to generate biomass and in particular mycolipids. For example, we show that Mtb re-routes carbon flux through a reversed MCC when growing on glycerol and Tween80. Finally, we find no evidence for PPDK flux under the conditions where it would be expected, indicating that Rv1127c is incorrectly annotated and that its actual role in the MCC remains unknown. Overall this work illustrates the flexibility of Mtb metabolism, and how it is able to adapt to the different combinations of nutrients that this pathogen might encounter during infection. The work also highlights the importance of exploring metabolic flux profiles in different conditions if we wish to fully comprehend metabolic potential of Mtb.

## Materials and Methods

### Bacterial strains and growth conditions

*Mycobacterium tuberculosis* (H37Rv) was grown in a 2 litre bioreactor (Adaptive Biosystem Voyager) under aerobic conditions and at pH 6.6 as previously described (4). Chemostat cultures were grown in Roisin’s minimal medium at a constant dilution rate of ~ 0.001 h^-1^ for cholesterol-acetate and 0.008 h^-1^ for glycerol-tween80 substrates respectively (slow growth rate). Culture samples were withdrawn from the chemostat to monitor cellular dry weight, viable counts, optical density, nutrient utilization and CO2 and O2 levels were measured in the exhaust gas (4). Metabolic steady state was confirmed from the OD600, substrate consumption and CO2 production rates.

Once metabolic steady state was reached ^13^C labelling experiments were initiated in the chemostat by replacing the feed medium with an identical medium containing a mixture of unlabelled and labelled carbon sources. Two mixtures of (i) 100% U-[^13^C_2_] acetate and unlabelled cholesterol or (ii) 30% U-[^13^C_3_] glycerol and unlabelled Tween80 were used as carbon substrates in Roisin’s minimal media. To confirm isotopic steady state, cultures were withdrawn at regular intervals and the amount of ^13^C label incorporation was measured using gas-chromatography mass spectrometry (GC-MS) analysis for every volume change.

To explore the potential flux through PPDK Mtb was grown in Roisin’s minimal medium containing 0.1 % (vol/vol) [3, 4-^13^C_2_] cholesterol as the sole carbon source until mid-log phase (OD_600_ = 0.6-0.8) before being harvested to analyse isotopologue incorporation into intracellular metabolites. To evaluate the incorporation of ^2^H into the amino acids of Mtb, deuterated cholesterol (0.0625 gL^-1^) was used as the single carbon source in Roisin’s media. Deuterated cholesterol was prepared using a recombinant Pichia pastoris strain CBS7435 Δhis4Δku70 Δerg5::pPpGAP-ZeocinTM-[DHCR7] Δerg6::pGAP-G418[DHCR24] (54). Cultures were adapted to growth in deuterated minimal in the Deuteration Laboratory (D-Lab) within the Life Sciences of the Institut Laue-Langevin (55) and purified as described by Moulin et al. (56). Mtb cultures grown in deuterated cholesterol were harvested at the mid-log phase for metabolomics.

### Culture analysis and substrate uptake measurements

Biomass and supernatant samples were collected and harvested (4). The amounts of glycerol in the supernatant and in fresh medium were assayed by use of a commercial assay kit that employs a glycerokinase-coupled enzyme assay system (Boehringer Mannheim). To assay for Tween 80 supernatant samples were hydrolysed by boiling for 1 h in methanolic KOH (5% potassium hydroxide: 50% methanol). After neutralizing the samples with HCl the free fatty acids were assayed using a commercial kit (Roche). Concentrations of cholesterol and acetate were measured using commercial kits supplied by Boehringer Mannheim.

### Biomass hydrolysate preparation and gas-chromatography mass-spectrometry (GC-MS) analysis

Biomass samples from the ^13^C labelled chemostat and mid-log batch cultures were washed and hydrolysed in 6M Hydrochloric acid for 24 hours (6). The dried hydrolysate was dissolved in 1 ml norvaline solution (0.075 mM in 80:20 H2O:MeOH). 0.1 ml was dried in-vacuo and this mixture was derivatised by adding 140μl acetonitrile:N-tert-butyldimethylsilyl-N-methyltrifluoroacetamide (MTBSTFA):1% tert butyldimethylchlorosilane (TBDMCSI), 1:1., sonicating (room temperature, 30 min) and then heating (90°C, 30 min) to complete the derivatisation. Samples were analysed by GC-MS within 72 h after derivatisation.

The analysis was performed with a GC system (Agilent) fitted with a DB-5ms capillary column (15 m × 0.18 mm internal diameter × 0.18 μm film with 5 m Duraguard integrated guard column) and deactivated quartz wool packed FocusLiner (SGE) coupled to a Pegasus III time-of-flight (TOF) mass spectrometer (Leco) equipped with a splitless injector. The injector temperature was initially held at 70°C for 2 min followed by heating to 350 °C at a rate of 17 °C min^-1^. This temperature was held for 1.5 min. The He Carrier flow was held constant at 1.4 ml min^-1^. The injection volume was 5 μl, inlet temperature was 250 °C, interface temperature at 310 °C and source temperature at 245°C. The system was operated with a mass range of 40–800 m/z at an EM voltage of 70 eV with a spectral acquisition rate of 20 spectra s^-1^. ^13^C isotopologue abundances (i.e., ^13^C incorporation; U-^12^C, ^13^C1,…, ^13^Cn) for each amino acid were determined for fragments containing the intact carbon skeleton for each amino acid; generally using the [M-57]^+^ ion. To obtain further information relating to the ^13^C incorporation at individual positions within individual amino acids, isotopologue abundances were also determined for fragments formed by cleavage of the C1–C2 bond in some samples. Within a subset of these, isotopologue abundances for the fragment formed by cleavage of the C2–C3 bond in serine were also determined allowing comprehensive characterization of the isotopic incorporation into serine. Mass spectra of the derivatized amino acids were corrected for the natural abundance of all stable isotopes (4, 6).

### Sample preparation and liquid-chromatography mass-spectrometry (LC-MS) analysis

For the detection of intracellular metabolites cultures were quenched in a solution of 60% methanol and 0.9% NaCl at −20 °C. The metabolites were extracted in a pre-chilled (−20°C) solution of methanol:chloroform (1:1) and cells were lysed twice by bead beating in the FastPrep at 6.5 for 20 seconds with careful cooling in between. Soluble extracts were filtered twice and then stored at −80°C until analysis by LC-MS.

Samples for ^13^C isotopomer profiling of organic acids and sugar phosphates were separated using a synergy hydro C18 reversed phase column on an Agilent 1200 chromatography system (Agilent Technologies, Waldbronn, Germany). Free amino acids were separated using a Luna SCX cation exchange column on a JASCO HPLC system. In both cases the HPLC outlets were coupled to an API 4000 (Applied Biosystems, Concord, Canada) mass spectrometer equipped with a TurboIon spray source. For detailed information on separation methods, optimal MS parameter settings as well as extraction and integration of ion chromatograms (58).

### MALDI lipid fingerprinting

A late log phase culture was washed in phosphate buffer saline by centrifugations and heat killed. The heat-inactivated was further washed three times in water and resuspended in 100 μl, An aliquot of 0.4 μl of the mycobacterial solution was loaded onto the target and immediately overlaid with 0.8 μl of 10 mg ml^-1^ super-2,5-dihydroxybenzoic acid (sDHB) matrix in chloroform/methanol (CHCl3/MeOH) 90:10 v/v. The mycobacterial sample and matrix were mixed directly on the target by pipetting and allowed to dry under a stream of air. MALDI-TOF MS analysis was performed on a 4800 Proteomics Analyzer (Applied Biosystems) using the reflectron mode. Samples were analysed by operating at 20 kV in the negative ion mode using an extraction delay time set at 20 ns.

### Metabolic modelling

Metabolic modelling was conducted using INCA, version 1.7 (59). An extended isotopomer model of central metabolism (Table S1) in Mtb was constructed of the model previously published in Beste et al., 2011. In addition to the reactions of glycolysis (EMP), gluconeogenesis, the pentose phosphate pathway (PPP), the tricarboxylic acid cycle (TCA) and anaplerosis (ANA), we included the reactions of the MCC and amino acid degradation pathways. This metabolic network model was supplemented with carbon atom transitions and consisted of a total of 90 reactions and 51 exchange reactions. Labelling measurements were obtained using 100% [U-^13^C_2_] acetate for CHL-ACE and 30% [U-^13^C_3_] glycerol for GLY-OLA. Flux values for net and exchange rates was derived from 86 independent flux parameters that are estimated using 108 labelling measurements for CHL-ACE and 120 for GLY-OLA respectively (supplementary data file S1). Substrate consumption rates for cholesterol, acetate, glycerol and Tween80 and overall CO2 net production rate (Table 1) were used as fixed fluxes to constrain the model.

### Metabolic flux estimation

Flux estimations were performed using INCA (59) using non-linear weighted least squares fitting approach to determine the flux values which are the most likely description of the labeling data and biomass constraints (4). INCA performs a levenberg-marquardt gradient-based search algorithm to minimise the sum of squared residuals (SSR) between the simulated and experimental measurements (27, 59). A multistart optimization approach was used for flux estimations with a total number of restarts of 100. Flux distributions with a minimum statistically acceptable SSR were considered the best-fit. The goodness-of-fit of the flux maps was asses by comparing the simulated and experimental measurements. The upper and lower 95% confidence limits of the best-fit flux distributions were calculated using parameter optimization option in INCA (59). Fluxes were considered to be significantly different if the 95% confidence limits do not overlap.

## Supplementary Figures

**Fig. S1.**
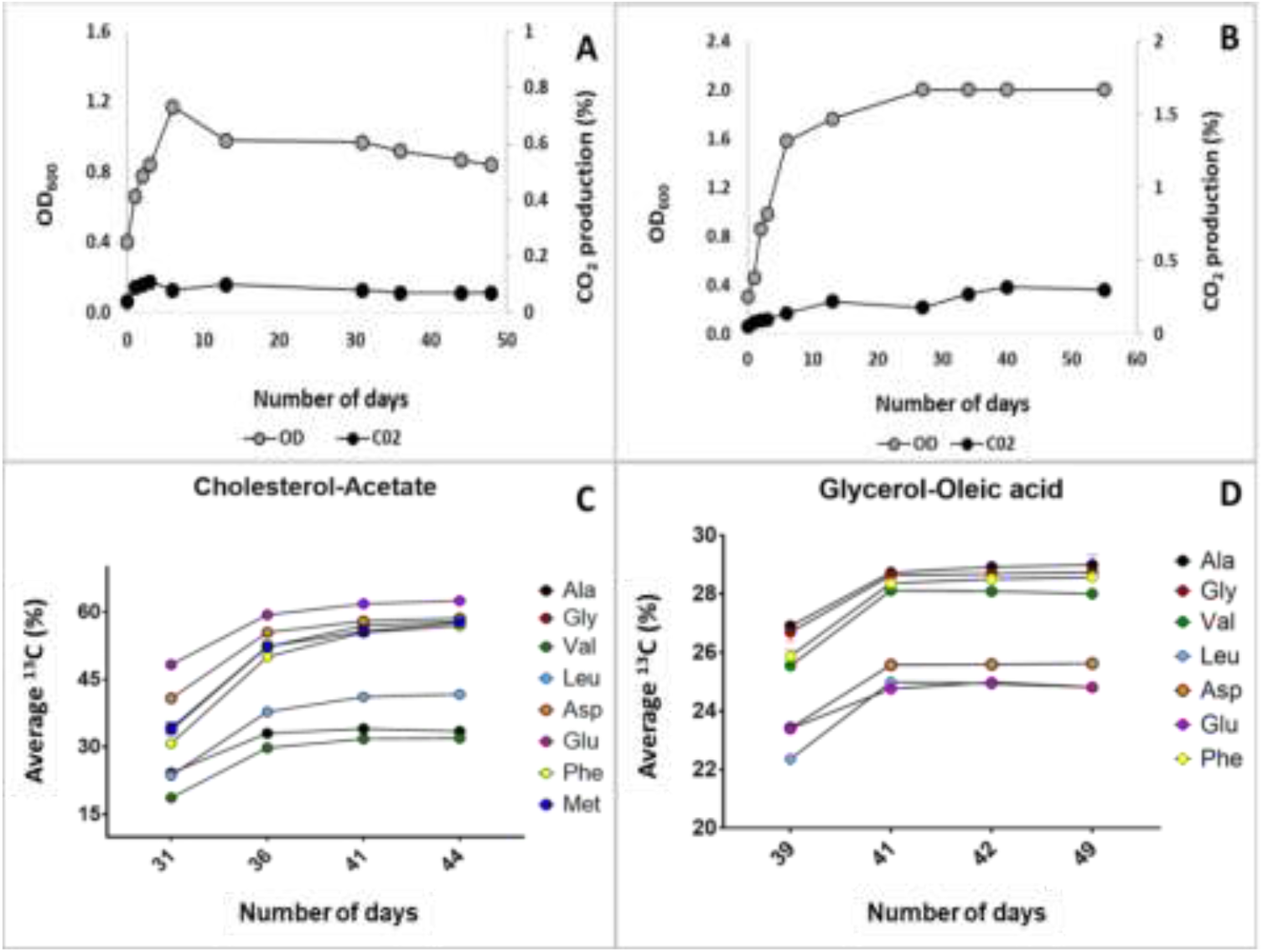
Metabolic and isotopic steady state growth of *Mycobacterium tuberculosis*. Continuous cultures of Mtb were grown in bioreactors with either CHL-ACE or GLY-OLA acid as carbon sources at a dilution rate of 0.01h^-1^ (t_d_ = 69 h). Metabolic steady state was achieved after three volume changes as evidenced from by stable cell density (OD_600_) and CO_2_ production rates (A-B). After metabolic steady state was achieved the feed was replaced with identical media containing ^13^C labelled carbon substrates until the culture attained an isotopic steady state as evidenced by steady levels of ^13^C incorporation into proteinogenic amino acids.

**Supplementary Fig. S2.**
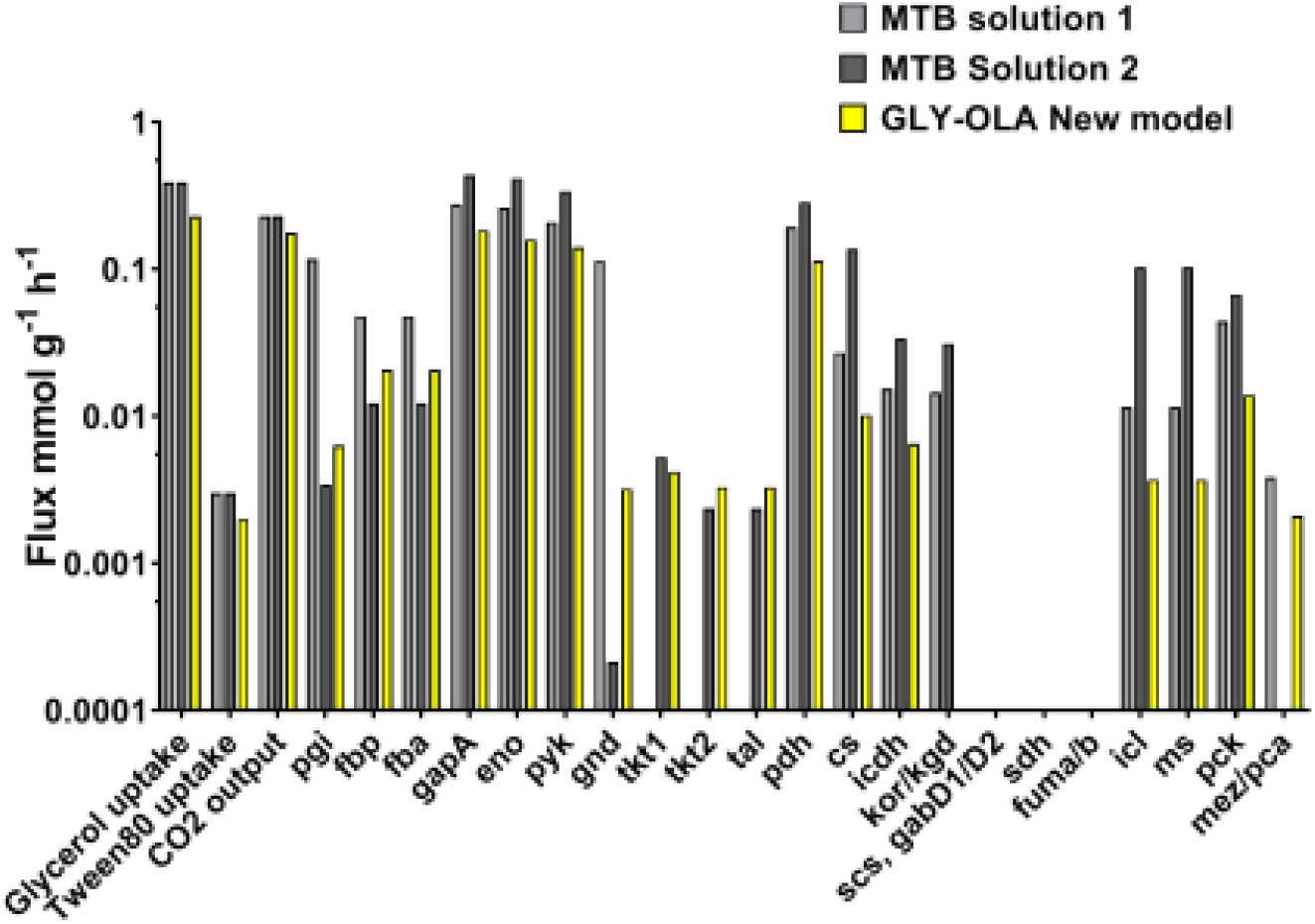
Absolute flux distributions. Fluxes measured in this study with the extended metabolic model was compared to the previously measured fluxes in Beste et al., 2011.

**Fig. S3.**
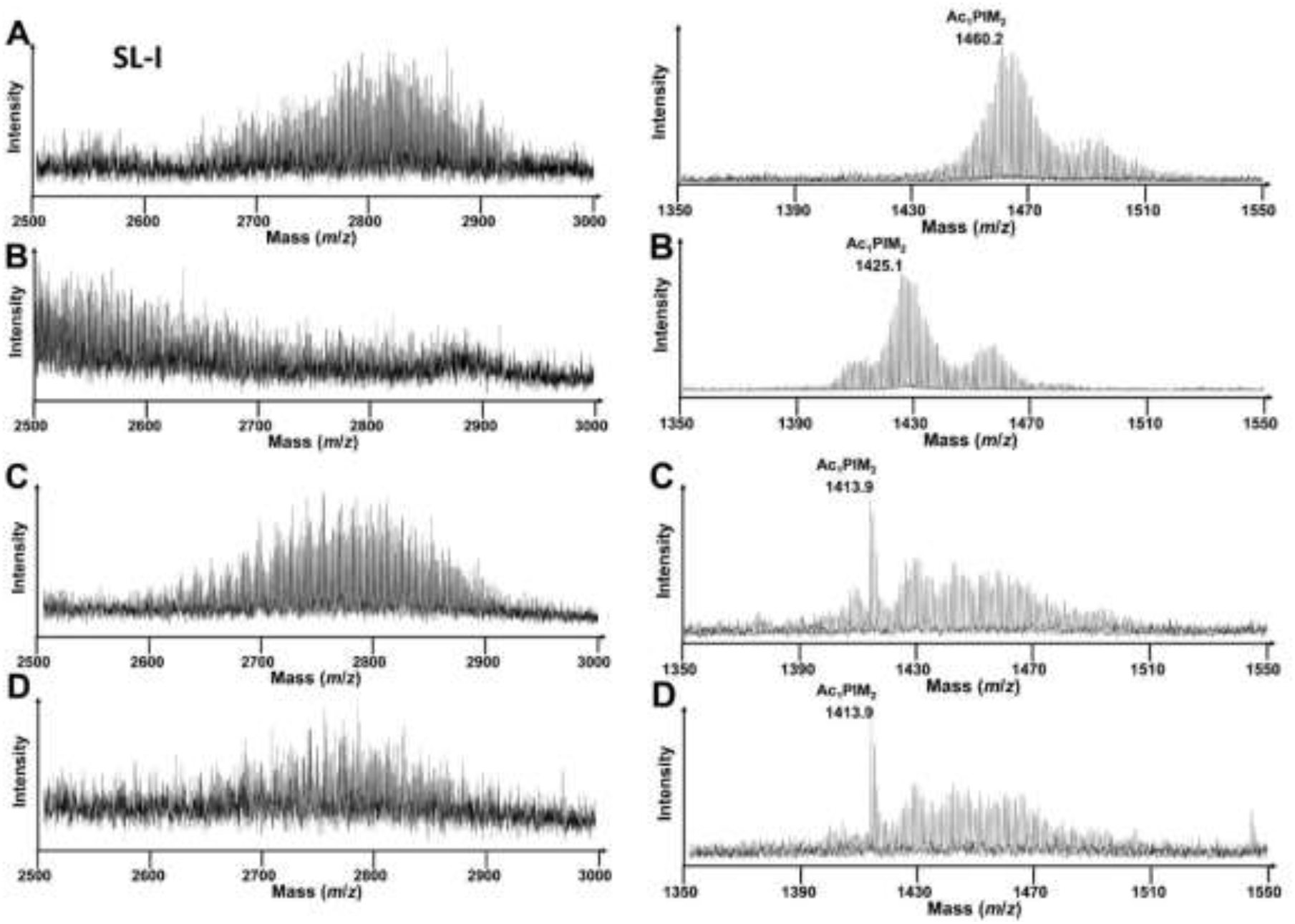
SL-I and Ac1PIM2 lipid finger printing. SL-I is observed as a broad set of peaks representing multiple lipoforms corresponding to differing numbers of CH_2_ units. Multiple lipoforms of SL-1 (m/z 2500 to m/z 3000) are also indicated SL-I. Mass shifts due to ^13^C labelling of whole bacteria lipid fingerprint prepared from metabolic and isotopic steady state chemostat for Mtb grown with [^13^C_2_] acetate and unlabelled cholesterol (A), [^13^C_3_]-glycerol and unlabelled Tween80 (B), unlabelled acetate and cholesterol (C) and unlabelled glycerol and Twen80 (D). Samples taken from the same chemostat at metabolic steady state (A) and (B) and isotopic in stationary state (C) and (D). The resolved mass shifts observed for Ac1PIM2 are shown to indicate significant lipid changes between the four conditions.

**Fig. S4.**
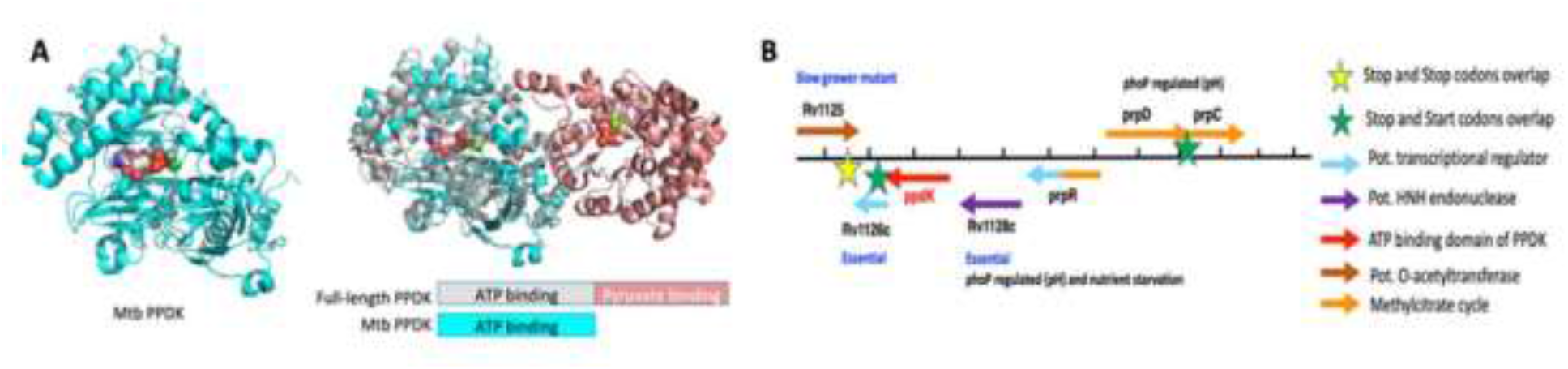
PPDK structure and genetic organisation in Mtb. (A) Left panel: iTasser model of Rv1127c (cyan) with ATP bound. Right panel: Overlay of the full-length PPDK (N-term domain white, C-term salmon with Rv1127c (cyan) showing that Rv1127c is lacking the PEP/pyruvate binding domain. (B) Mtb genomic organization of Rv1127c (ppdk, red arrow).

**Table S1.**
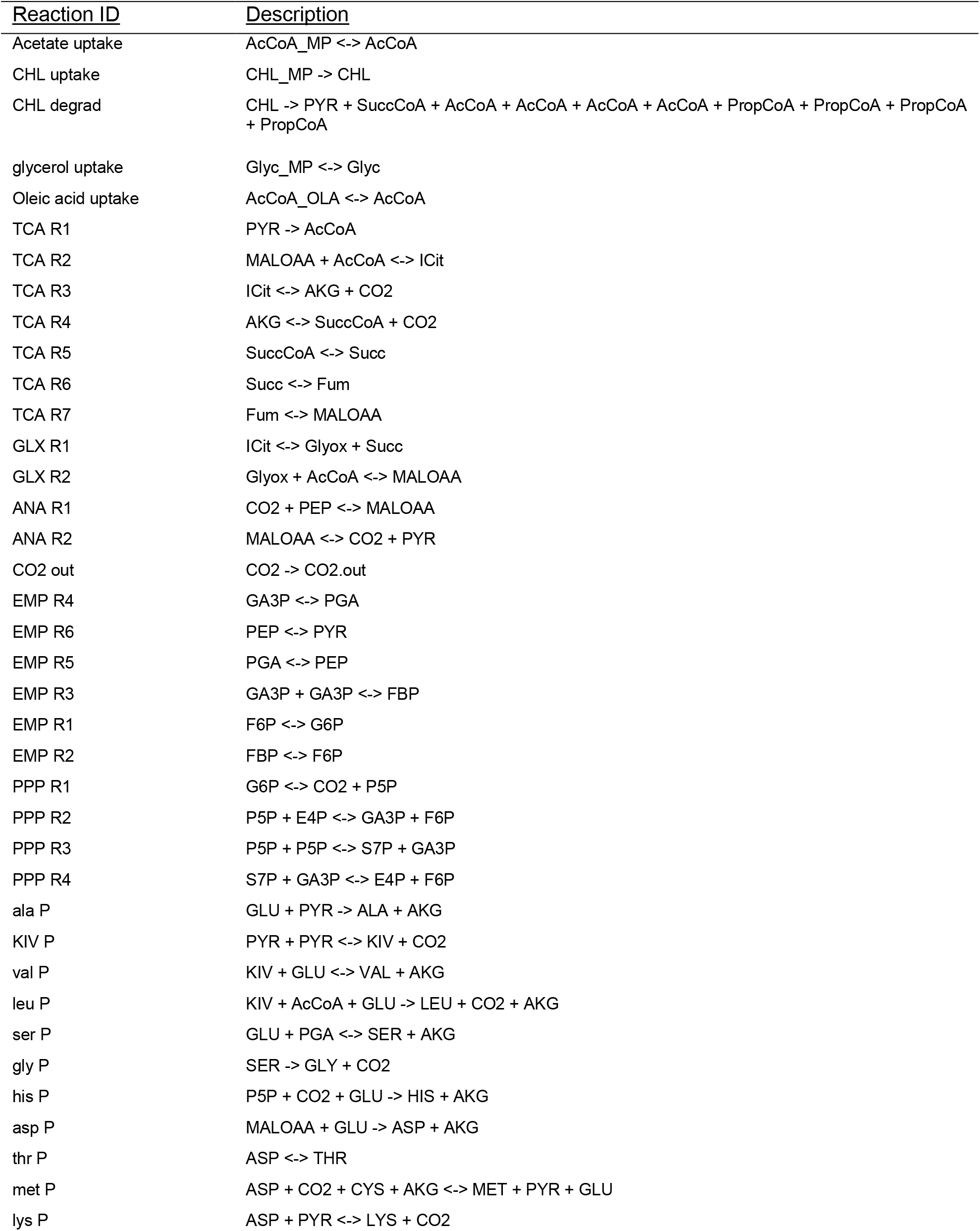
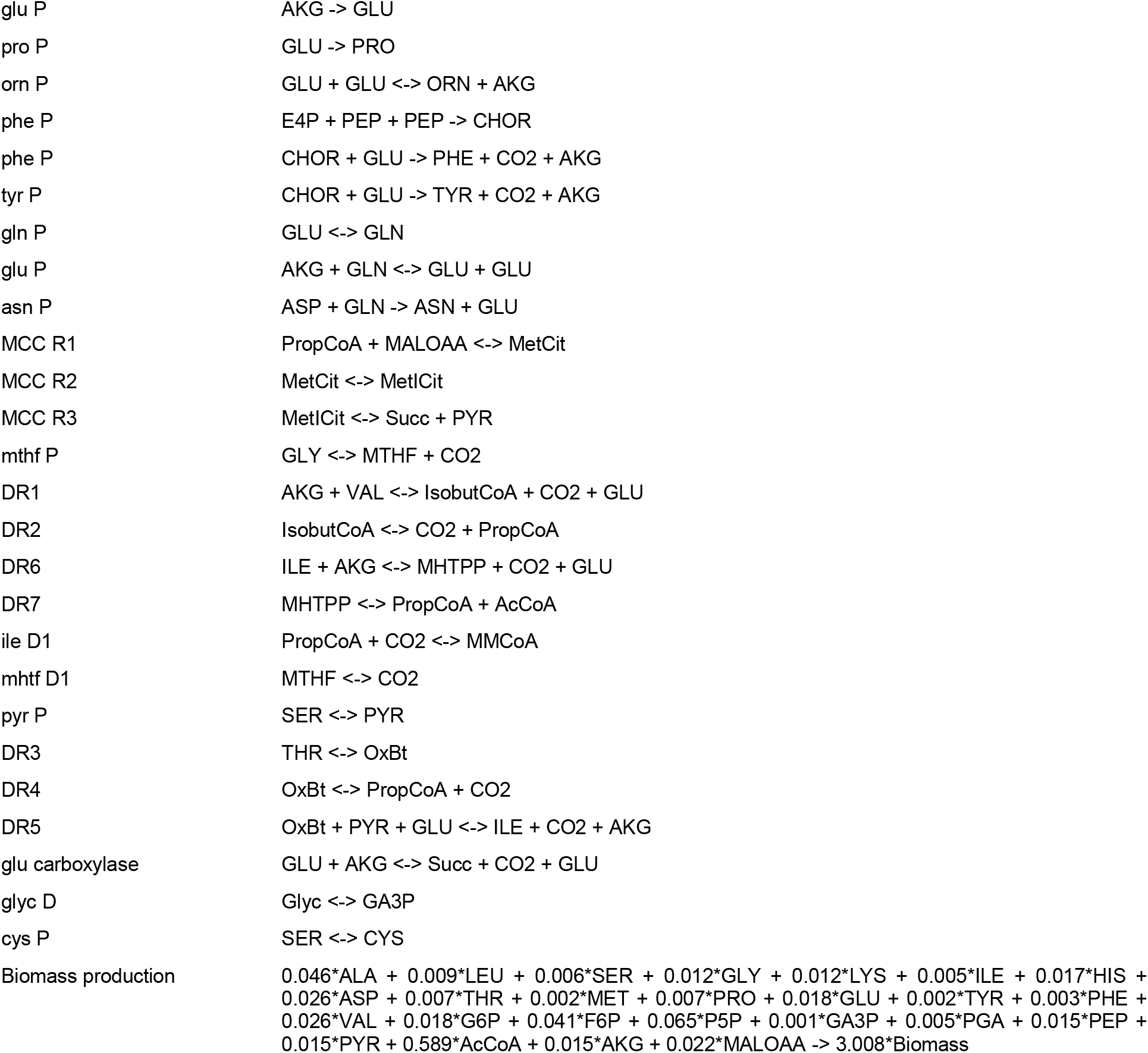

| Reaction ID | Description |
| --- | --- |
| Acetate uptake | AcCoA_MP <-> AcCoA |
| CHL uptake | CHL_MP -> CHL |
| CHL degrad | CHL -> PYR + SuccCoA + AcCoA + AcCoA + AcCoA + AcCoA + PropCoA + PropCoA + PropCoA + PropCoA |
| glycerol uptake | Glyc_MP <-> Glyc |
| Oleic acid uptake | AcCoA_OLA <-> AcCoA |
| TCA R1 | PYR -> AcCoA |
| TCA R2 | MALOA + AcCoA <-> ICit |
| TCA R3 | ICit <-> AKG + CO2 |
| TCA R4 | AKG <-> SuccCoA + CO2 |
| TCA R5 | SuccCoA <-> Succ |
| TCA R6 | Succ <-> Fum |
| TCA R7 | Fum <-> MALOA |
| GLX R1 | ICit <-> Glyox + Succ |
| GLX R2 | Glyox + AcCoA <-> MALOA |
| ANA R1 | CO2 + PEP <-> MALOA |
| ANA R2 | MALOA <-> CO2 + PYR |
| CO2 out | CO2 -> CO2.out |
| EMP R4 | GA3P <-> PGA |
| EMP R6 | PEP <-> PYR |
| EMP R5 | PGA <-> PEP |
| EMP R3 | GA3P + GA3P <-> FBP |
| EMP R1 | F6P <-> G6P |
| EMP R2 | FBP <-> F6P |
| PPP R1 | G6P <-> CO2 + P5P |
| PPP R2 | P5P + E4P <-> GA3P + F6P |
| PPP R3 | P5P + P5P <-> S7P + GA3P |
| PPP R4 | S7P + GA3P <-> E4P + F6P |
| ala P | GLU + PYR -> ALA + AKG |
| KIV P | PYR + PYR <-> KIV + CO2 |
| val P | KIV + GLU <-> VAL + AKG |
| leu P | KIV + AcCoA + GLU -> LEU + CO2 + AKG |
| ser P | GLU + PGA <-> SER + AKG |
| gly P | SER -> GLY + CO2 |
| his P | P5P + CO2 + GLU -> HIS + AKG |
| asp P | MALOA + GLU -> ASP + AKG |
| thr P | ASP <-> THR |
| met P | ASP + CO2 + CYS + AKG <-> MET + PYR + GLU |
| lys P | ASP + PYR <-> LYS + CO2 |
| glu P | AKG -> GLU |
| pro P | GLU -> PRO |
| orn P | GLU + GLU <=> ORN + AKG |
| phe P | E4P + PEP + PEP -> CHOR |
| phe P | CHOR + GLU -> PHE + CO2 + AKG |
| tyr P | CHOR + GLU -> TYR + CO2 + AKG |
| gln P | GLU <=> GLN |
| glu P | AKG + GLN <=> GLU + GLU |
| asn P | ASP + GLN -> ASN + GLU |
| MCC R1 | PropCoA + MALOAA <=> MetCit |
| MCC R2 | MetCit <=> MetICit |
| MCC R3 | MetICit <=> Succ + PYR |
| mthf P | GLY <=> MTHF + CO2 |
| DR1 | AKG + VAL <=> IsobutCoA + CO2 + GLU |
| DR2 | IsobutCoA <=> CO2 + PropCoA |
| DR6 | ILE + AKG <=> MHTPP + CO2 + GLU |
| DR7 | MHTPP <=> PropCoA + AcCoA |
| ile D1 | PropCoA + CO2 <=> MMCoA |
| mhtf D1 | MTHF <=> CO2 |
| pyr P | SER <=> PYR |
| DR3 | THR <=> OxBt |
| DR4 | OxBt <=> PropCoA + CO2 |
| DR5 | OxBt + PYR + GLU <=> ILE + CO2 + AKG |
| glu carboxylase | GLU + AKG <=> Succ + CO2 + GLU |
| glyc D | Glyc <=> GA3P |
| cys P | SER <=> CYS |
| Biomass production | 0.046*ALA + 0.009*LEU + 0.006*SER + 0.012*GLY + 0.012*LYS + 0.005*ILE + 0.017*HIS + 0.026*ASP + 0.007*THR + 0.002*MET + 0.007*PRO + 0.018*GLU + 0.002*TYR + 0.003*PHE + 0.026*VAL + 0.018*G6P + 0.041*F6P + 0.065*P5P + 0.001*GA3P + 0.005*PGA + 0.015*PEP + 0.015*PYR + 0.589*AcCoA + 0.015*AKG + 0.022*MALOAA -> 3.008*Biomass |

